# Overwriting an instinct: visual cortex instructs learning to suppress fear responses

**DOI:** 10.1101/2024.07.31.605567

**Authors:** Sara Mederos, Nicole Vissers, Patty Blakely, Claudia Clopath, Sonja Hofer

## Abstract

Fast instinctive responses to environmental stimuli can be crucial for survival, but are not always optimal. Based on prior experience, animals can thus adapt their behavior and suppress instinctive reactions. However, the neural pathways mediating such ethologically relevant forms of learning remain unclear. We show that posterolateral higher visual areas (plHVAs) are crucial for learning to suppress escapes from innate visual threats through a top-down pathway involving the ventrolateral geniculate nucleus (vLGN). plHVAs are no longer necessary after learning: instead, the learnt behavior relies on plasticity within vLGN populations that exert inhibitory control over fear responses. vLGN neurons receiving input from plHVAs enhance their responses to visual threat stimuli during learning through endocannabinoid-mediated long-term suppression of their inhibitory inputs. We thus reveal the detailed circuit, cellular and synaptic mechanisms underlying experience-dependent suppression of fear responses through a novel corticofugal pathway.

## Introduction

Instinctive behaviors are automatic responses to specific environmental challenges that have evolved to furnish animals with a repertoire of behaviors vital for survival and reproductive success. These behaviors allow animals to quickly detect and respond to potential dangers or opportunities in their environment without the need for prior learning or experience (*1*, *2*), and are usually implemented by brainstem pathways independent of neural processes in the forebrain (*3*– *5*). However, to ensure continuing success in changing environments, it is also important to be able to suppress instinctive reactions if they are no longer appropriate or advantageous (*6–9*). And indeed, many animals have the capacity to modify instinctive behaviors based on experience or changing circumstances (*5*, *7*, *10–14*). This behavioral flexibility allows animals to fine-tune their responses to their specific environment and can serve to conserve resources, avoid unnecessary risks, or capitalize on new opportunities. However, the neural basis of this ethologically highly relevant form of learning, the overwriting of instinctive reactions, is still unclear. In this study we test the hypothesis that neocortical circuits can regulate brainstem-driven instincts, enabling flexible behavior.

Fear responses to visual threats, such as escape from an approaching aerial predator, are examples of instinctive reactions particularly crucial for survival (*1*, *3*, *4*, *15*, *16*). Escapes from overhead looming stimuli mimicking aerial predators are mediated by neural circuits involving the medial superior colliculus and the periaqueductal gray (*3*, *17–19*). This visuo-motor pathway in the brainstem can autonomously drive escape responses independently of the forebrain (*3*, *20*). However, animals can suppress these fear responses as they learn that a perceived visual threat proves harmless (*5*, *7*, *10*, *11*, *14*, *21*), and this form of adaptive behavior may involve neocortical circuits. Sensory circuits in the neocortex have previously been shown to modulate different forms of instinctive or reflexive reactions to sensory stimuli (*12*, *22*, *23*). In particular, higher visual areas (HVAs) in rodents can integrate both visual and diverse behavioral and task-related variables and have been linked to numerous functions that go beyond basic visual processing (*24–28*). HVAs posterolateral to the primary visual cortex (V1) contribute to learning and execution of various learned visually-guided behaviors, are modified by prior experience and have been shown to adaptively modulate innate behaviors (*22*, *24*, *25*, *29–31*). These areas, including postrhinal (POR), lateromedial (LM), posteriomedial (P) and laterointermediate (LI) cortex, which we will collectively refer to as posterolateral HVAs (plHVAs), thus may provide suitable candidate regions for implementing experience-dependent control over visually-driven instincts through their extensive cortico-fugal projections. One pathway that provides visual cortical areas with a route to exert strong inhibitory control over brainstem processing and thus over behavioral output, is the dense projection to the ventral lateral geniculate nucleus (vLGN) in the prethalamus (*11*, *32–34*). Prethalamic areas including the vLGN and the adjacent zona incerta are part of the diencephalon, consist mainly of GABAergic neurons and have been shown to act as inhibitory control hubs of diverse instinctive behaviors (*33*, *35*). The vLGN in particular receives visual input from the retina and has powerful control over fear responses to visual threat (*11*, *33*, *34*). Here we investigate if and how the corticofugal pathway from plHVAs to vLGN mediates learned suppression of fear responses, and can thus provide a cognitive control mechanism enabling adaptive behavior shaped by experience.

## Results

Escape behavior evoked by a looming (i.e. dark overhead expanding) stimulus is a well-established protocol for assessing instinctive fear responses (*3*, *15*). When naïve mice are presented with this visual stimulus, they consistently escape to a shelter provided at the other end of an elongated arena (Movie 1). However, mice can adapt their behavior and suppress this fear response if they learn that the potential threat stimulus does not result in negative consequences (*10*, *11*, *14*). However, if animals are given the opportunity to seek shelter, they often keep escaping to high-contrast looming stimuli for many stimulus repetitions and several behavioral sessions (Fig. S1A-D). We therefore adapted a protocol developed by Lenzi and colleagues (*10*), in which we prevented access to the shelter with a dividing barrier and presented looming stimuli with increasing contrast (acquisition phase, Fig. 1A and Movie 2). We then removed the barrier and assessed the likelihood of mice to escape from high-contrast looming stimuli (probe phase, Fig. 1A). Control mice showed strongly decreased fear responses after this learning protocol, and only rarely escaped from the looming stimulus (Fig. S1E, Movie 3).

**Figure 1.**
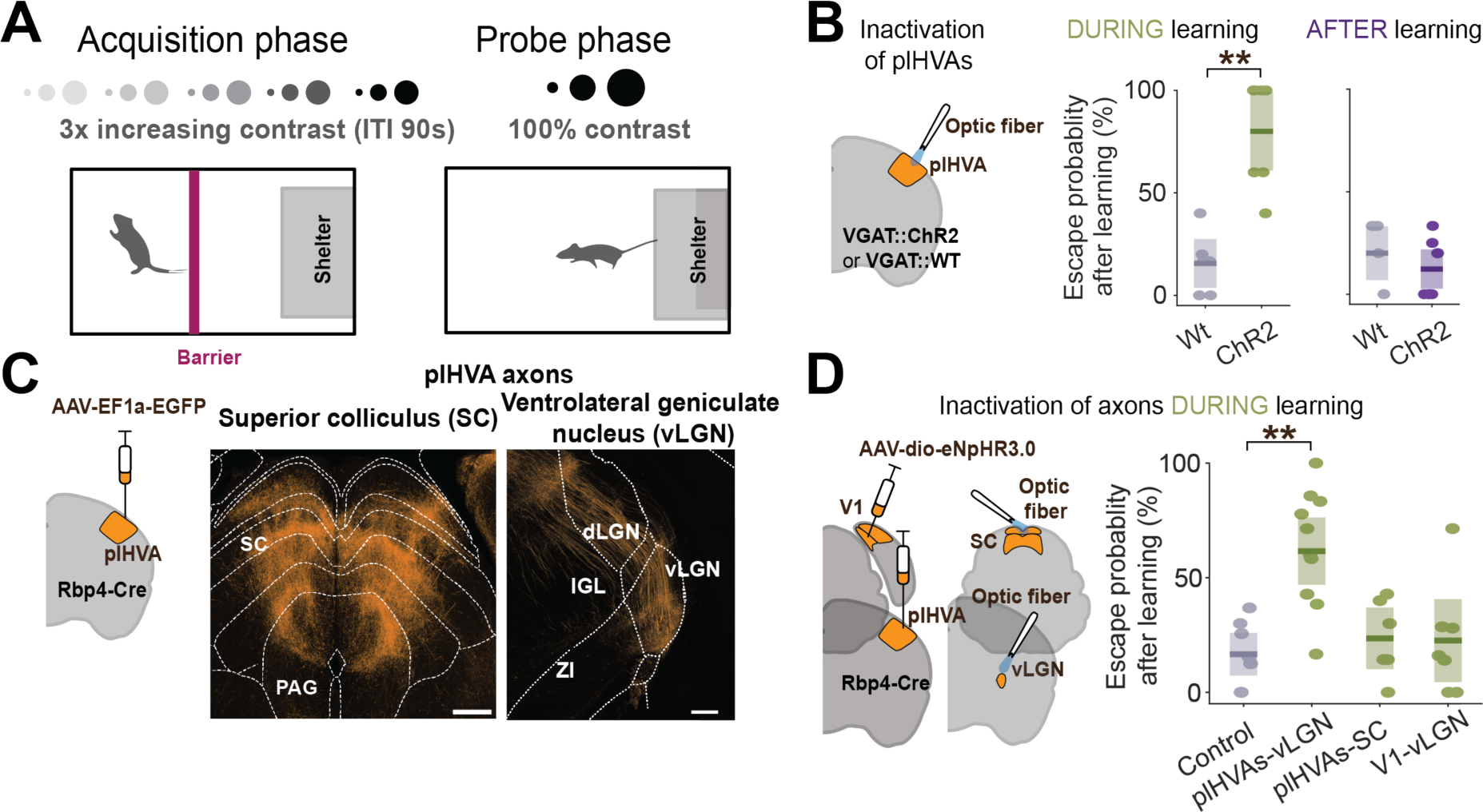
plHVAs instruct learning through corticofugal projections to the ventral lateral geniculate nucleus (vLGN) **(A)** Schematic of the task and stimulus protocols used to elicit learning to suppress escape. Acquisition phase: a barrier is used to prevent escape to a shelter while looming stimuli with increasing contrast are shown sequentially. ITI: inter trial interval. Probe phase: the barrier is removed and 6 to 10 100%-contrast looming stimuli are shown. **(B)** Left, experimental approach. Middle, boxplot (showing median and interquartile range) of escape probability after learning with (green) and without (grey) silencing of plHVAs during learning. Dots show individual animals. P = 0.003, KW test; n = 6 (wildtype control (wt)) and 7 mice (ChR2). Right, same as middle but plHVAs are silenced only after learning. P = 0.222, KW test; n = 6 (control) and 8 mice (ChR2, purple). **(C)** Left, schematic of tracer injection in plHVAs. Right, labeled plHVA axons in different target areas. Scale bars 500 μm, 250 μm. PAG: periaqueductal gray, SC: superior colliculus, vLGN: ventral lateral geniculate nucleus, IGL: Intergeniculate leaflet, dLGN: dorsal geniculate nucleus, ZI: zona incerta. **(D)** Experimental design. Right, boxplots of escape probabilities after learning in control animals and when plHVA axons in vLGN (plHVA-vLGN) or SC (plHVA-SC) are silenced, or when V1 axons in vLGN are silenced during learning (V1-vLGN). Dots show individual animals. plHVA-vLGN vs control: P = 0.003; plHVA-SC vs control: P = 0.856; V1-vLGN vs control: P = 0.973; KW test; n = 8, 11, 6 and 7 mice. Statistical significance is indicated as ** P < 0.01.

To test if neural activity in plHVAs is important for this learned suppression of escape responses, we used transgenic mice expressing channelrhodopsin-2 (ChR2) in GABAergic neurons and implanted optical fibers to silence neural activity bilaterally in plHVAs with blue light (*36*). Silencing plHVAs while presenting looming stimuli in naïve mice had no effect on the animals’ probability to escape (Fig. S1F), consistent with previous results showing that visual cortex has little influence on the instinctive escape response to visual threats (*3*). However, when plHVAs were silenced during the learning protocol (during all stimulus presentations in the acquisition phase), mice failed to learn and still showed high escape probabilities to looming stimuli in the probe phase (Fig. 1B, left). We repeated these experiments with a long learning protocol with access to the shelter, in which mice eventually ceased to escape after presentation of many high-contrast looming stimuli (over several days, Fig. S1A-D). Silencing plHVAs using muscimol had little effect on instinctive escape behavior in naïve mice, but, when applied throughout the learning protocol, prevented mice from learning to suppress escape responses (Fig. S1A-D) corroborating our results of silencing plHVAs in the shorter learning protocol. plHVAs are thus necessary for learning to suppress escape responses independent of the specific experimental protocol used to induce learning. In contrast, optogenetic silencing of plHVAs during the probe phase, i.e. only after mice had learnt, had no effect on the learnt behavior, as mice still suppressed escape responses when plHVAs were silenced (Figure 1B, right). This indicates that neural activity in plHVAs is crucial for learning to suppress escape behavior but is not necessary to execute and sustain this learnt behavioral response after learning.

We next aimed to identify the specific pathways through which plHVAs mediate learnt suppression of fear responses. Anterograde axon labeling from layer 5 (L5) neurons in plHVAs using Rbp4-Cre mice showed dense projections in several subcortical regions, including the medial and deep layers of the superior colliculus (SC), previously shown to be crucial for generating escape responses to looming stimuli (*3*, *17*). Another clear target of plHVA projections was the ventral lateral geniculate nucleus (vLGN) (Fig. 1C), an inhibitory prethalamic area that has strong control over fear behavior and which, when activated, can fully block escape responses by inhibiting SC activity (*11*, *33*, *34*). To test the relevance of these two plHVA pathways for learning, we optogenetically silenced plHVA axonal projections selectively in either SC or vLGN by optical stimulation of halorhodopsin (eNpHR3.0) expressing plHVA axons during the acquisition phase of the learning protocol (Fig. 1A, D). Silencing plHVA projections to vLGN prevented learning, as mice continued to escape to the looming stimulus afterwards (Fig. 1D). In contrast, silencing plHVA projections to SC had no effect on learning (Fig. 1D). Moreover, silencing projections from V1 to vLGN also did not affect learning, showing that it is specifically projections from plHVA to vLGN that are necessary for mice to learn to suppress escape responses (Fig. 1D & S2C, D). We corroborated the necessity of the plHVA->vLGN pathway for learning with a chemogenetic approach where we targeted the inhibitory designer receptor hM4Di selectively to plHVA L5 neurons in Rbp4-cre mice and applied the agonist CNO locally in vLGN (Fig. S2E, F). Silencing projections from plHVA to vLGN in this way also impeded learning, such that mice were still likely to escape from the looming stimulus after the learning protocol.

The experience of looming stimuli during the learning protocol likely induces lasting changes in neural circuits, i.e. a memory of this prior experience, leading to the adapted behavioral response to the visual threat stimulus. Our data show that while plHVAs are required for learning to suppress fear responses, they are no longer necessary after learning. This indicates that the learning-induced memory is stored in neural circuits downstream of plHVAs. Since activation of vLGN by plHVA projections is necessary for learning, vLGN is one potential substrate for memory formation. We therefore set out to test this hypothesis. First, we examined if activity in vLGN is required for the adapted behavioral response, i.e. the suppression of escape to looming stimuli after learning. We expressed Cre-dependent stGtACR2 in vLGN of VGAT-Cre mice via AAV injections to optogenetically silence inhibitory cells which constitute the large majority of vLGN neurons (*11*, *32*, *33*). When we silenced vLGN during stimulus presentation after learning (in the probe phase), mice resumed escaping from looming stimuli (Fig. 2A, B). This suggests that - unlike plHVAs - vLGN is necessary for the learned behavioral response. Mice immediately reverted to the learned behavior of suppressing escape responses when optogenetic manipulation was switched off, indicating that transient inhibition of vLGN neurons did not cause a sustained increase in fear or anxiety-related state (Fig. 2B).

**Figure 2.**
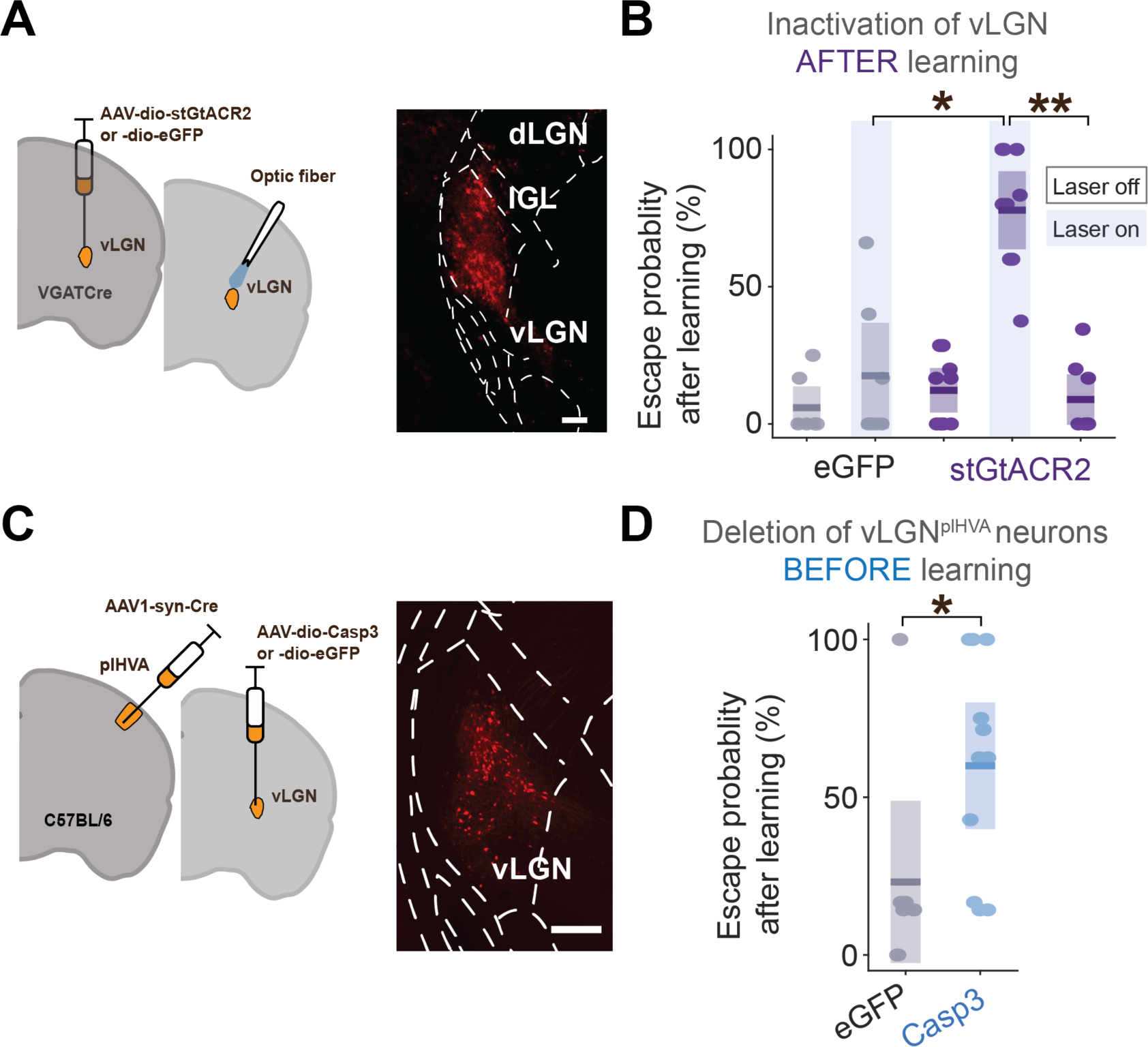
vLGN cells receiving plHVA input are necessary for learning. **(A)** Experimental approach for acute inhibition of vLGN. Right, example image of vLGN neurons expressing stGtACR2. Scale bar 150 μm. **(B)** Boxplot (showing median and IQR) comparing post-learning escape probabilities to 6 - 10 high-contrast looming stimuli for control mice expressing eGFP in vLGN with and without laser (gray, 7 mice, P = 0.954, KW test) and mice with stGtACR2 expression in GABAergic vLGN neurons without laser, with laser and subsequently without laser again (purple, 9 mice, without before vs with laser: P = 0.0057 repeated-measures ANOVA; with laser vs without laser after: P = 0.0020, repeated-measures ANOVA; Laser eGFP vs Laser stGtACR2 P = 0.005, KW test). **(C)** Left, Experimental approach for specifically lesioning vLGN cells receiving input from plHVAs. Right, Example image of eGFP-labeled vLGN cells. Scale bar 200 μm. **(D)** Boxplot of post-learning escape probabilities of control (gray, 7 mice) and mice with vLGN cells receiving input from plHVAs ablated before learning (blue, 11 mice, P = 0.043, KW test). Statistical significance is indicated as * P < 0.05. Dots represent individual animals.

However, silencing of vLGN could generally lower the threshold for threat-evoked escape responses independently of the plHVA-dependent learning process (*11*, *34*). To test the role of the plHVA-vLGN pathway in learned suppression of escape responses more specifically, we selectively targeted vLGN neurons receiving input from plHVAs by combining anterograde transfer of Cre-recombinase from plHVAs to vLGN and Cre-dependent gene expression in vLGN (see Methods). vLGN neurons receiving input from plHVAs were GABAergic cells projecting to SC and other target areas (Fig. S3). When we selectively ablated plHVA-innervated vLGN neurons using Cre-dependent caspase expression (Fig. 2C), mice showed impaired learning with a higher likelihood to escape from looming stimuli than control mice after the learning protocol (Fig. 2 C,D). This indicates that plHVA-innervated vLGN neurons are necessary for learning, and learning-induced neural changes may be expressed within vLGN or in its connections to downstream areas.

Our findings suggest that plHVA input drives learning by changing the activity of vLGN neurons able to suppress fear responses. To determine how neural activity in vLGN changes during learning we performed electrophysiological single-unit recordings during presentation of high-contrast looming stimuli. We recorded over many stimulus presentations until animals learned not to escape (long learning protocol, see also Fig. S1), using chronically implanted silicon probes in vLGN (Fig. 3A & Fig. S4A), and selected cells responsive to the looming stimulus within 0 to 300 ms after stimulus onset (before learning, after learning or both). Restricting the analysis to this early time window allowed us to isolate visual signals and minimize the influence of motor-related activity, because the average escape latency of mice was 1.73 ± 0.08 s (mean =/- std), and we excluded the few trials in which mice initiated an escape earlier than 300 ms after stimulus onset (Fig. S4B). vLGN neurons exhibited diverse responses to looming stimuli and many neurons changed their activity during learning. To capture such changes, we divided neurons depending on whether they significantly increased their firing rate, decreased their firing rate or showed no change in looming stimulus response over learning (Fig. 3B, C).

**Figure 3.**
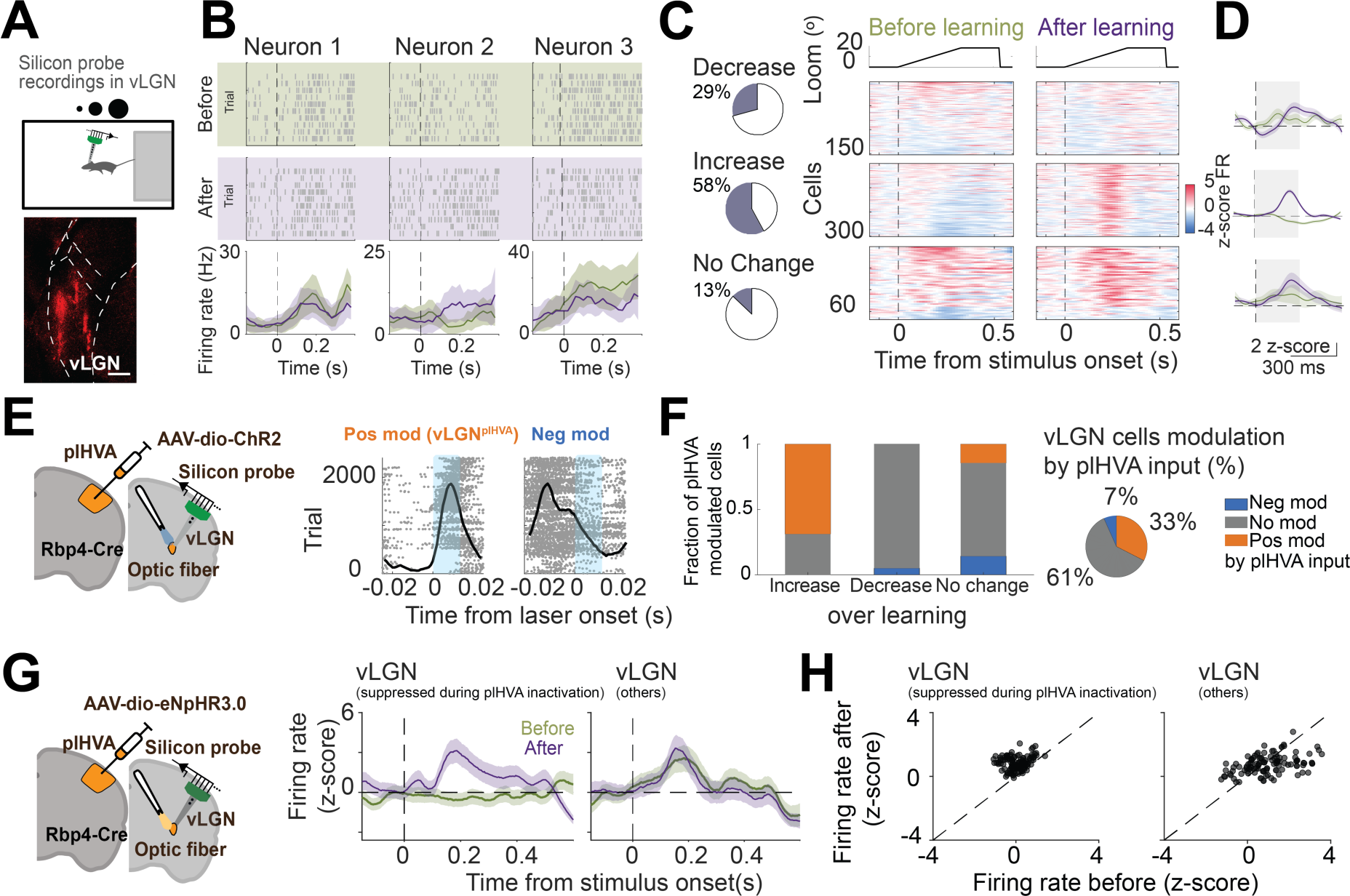
vLGN neurons receiving input from plHVAs increase their response to looming stimuli during learning. **(A)** Top, experimental approach: chronic electrophysiological recordings in vLGN while freely-moving animals are exposed to high-contrast looming stimuli until they learn to suppress escape responses. Bottom, example image with probe shank locations. Scale bar 200 μm. **(B)** Looming stimulus responses of 3 vLGN example cells before and after learning. Blue dots indicate escape onset. **(C)** Left, Pie charts of fraction of vLGN neurons with increased and decreased responses or no change in response strength over learning in a time window of 0 - 300 ms after stimulus onset. Right, z-scored spike rate responses aligned to looming stimulus onset of all isolated units classified as looming-stimulus responsive neurons (from n = 9 mice) before and after learning, allocated according to their response change over learning. **(D)** Left, mean peristimulus histograms (PSTHs) of looming responses before and after learning of the three groups of vLGN neurons in C. Dashed line shows stimulus onset, shaded area denotes time window used for analysis (n = 523; n = 285; n = 94 cells from n = 9 mice). **(E)** Left, Experimental design showing silicon probe placement combined with an optic fiber for stimulation of ChR2-expressing plHVA axons. Right, spike responses to optogenetic stimulation and PSTHs for example vLGN neurons. **(F)** Left, Fraction of negatively modulated (neg mod), positively modulated (pos mod), and non-modulated neurons (no mod) during stimulation of plHVAs for vLGN neurons that increase, decrease or show no change in response to looming stimuli over learning. Pie charts show the fraction of units exhibiting modulation by plHVA activation out of all recorded neurons (n = 175; n = 302; n = 36 cells from 5 mice). **(G)** Left, Experimental design. Electrophysiological recordings from vLGN cells with plHVA silencing during a subset of looming stimuli presentations. Right, Mean PSTHs in response to looming stimuli for neurons suppressed during plHVA silencing (left) and other vLGN neurons (right). (n=108; n=281 cells from 4 mice) **(H)** Scatterplots of looming stimulus response strength before and after learning (0 - 300 ms after stimulus onset) for individual vLGN neurons divided as in G (suppressed during plHVA silencing P < 0.0001; Wilcoxon Signed-Rank Test; and other vLGN neurons: P = 0.0521; Wilcoxon Signed-Rank Test).

In a subset of animals, we expressed ChR2 in L5 neurons of plHVAs in Rbp4-Cre mice to combine electrophysiological recordings of vLGN cells with optogenetic activation of plHVA axons in vLGN. This allowed us to identify vLGN neurons excitated by plHVAs (positively modulated), inhibited (negatively modulated) or not affected by plHVA axon activation, and these groups of neurons showed differences in their electrophysiological properties and in how their looming stimulus responses changed over learning (Fig. 3E, F, Fig. S4C-E). Strikingly, we found that the large majority of vLGN neurons excited by plHVAs increased their responses to looming stimuli during learning (Fig. 3E, F), indicating a crucial role for plHVA inputs in shaping vLGN responses to looming stimuli over learning. To corroborate these findings and investigate how plHVA inputs contribute to looming responses of vLGN neurons, we performed another set of experiments in which we optogenetically inactivated plHVAs inputs to vLGN using eNpHR3.0 during a subset of interleaved looming stimuli trials while recording from vLGN neurons. We identified vLGN neurons receiving excitatory input from plHVAs as those that were significantly suppressed during plHVA inactivation (Fig. S5A-C). These vLGN neurons again showed, on average, a clear increase in looming stimulus responses over learning (Fig. 3G, H), even when plHVA input was removed (Fig. S5A-C). In contrast, the remaining vLGN population showed no response change over learning (Fig. 3G, H and Fig. S5). Both plHVA activation and inactivation experiments are thus in strong concordance, showing that it is predominantly those vLGN neurons receiving excitatory input from plHVAs that change their neural activity during learning, resulting in an increased spiking response to looming stimuli. Importantly, as demonstrated above, these vLGN neurons are necessary for learning to suppress escape responses (Fig. 2C, D). Together, these results indicate that plHVA activity elicits plastic changes within vLGN circuits during learning, causing a persistent increase in looming-evoked responses of plHVA-driven vLGN neurons. Such increased activity of inhibitory vLGN neurons can then inhibit behavioral escape responses (*11*, *34*).

Next, we set out to explore the potential cellular and synaptic mechanisms of this learning-induced change in vLGN activity. To determine if functional interactions between different groups of vLGN neurons change during learning, we calculated pairwise cross-correlation functions of the spike trains of all simultaneously recorded vLGN neurons. Interestingly, neurons that were positively modulated by plHVA activation again stood out in that many of them had negative spike time correlations with the rest of the vLGN network, especially with the neurons not affected by plHVA activation (Fig. 4A-C, Fig. S6A, B). For many of these cell pairs, the troughs in the cross correlogram were biased towards positive time lags (Fig. S6C), suggesting that neurons that receive plHVA input may be inhibited by the local vLGN network before learning. This is consistent with the observation that these vLGN neurons decrease their activity in response to looming stimuli before learning, in particular when excitatory plHVA input is removed (Fig. S5C). Interestingly, spike timing relationships specifically between vLGN neurons positively modulated by plHVA activation and the rest of the network changed with learning, such that these neurons showed positive correlations with the remaining vLGN neurons after learning (Fig. 4A-C, Fig. S6). This suggests that inhibitory drive, potentially from within vLGN, onto vLGN neurons that receive direct input from plHVAs is decreased during learning.

**Figure 4.**
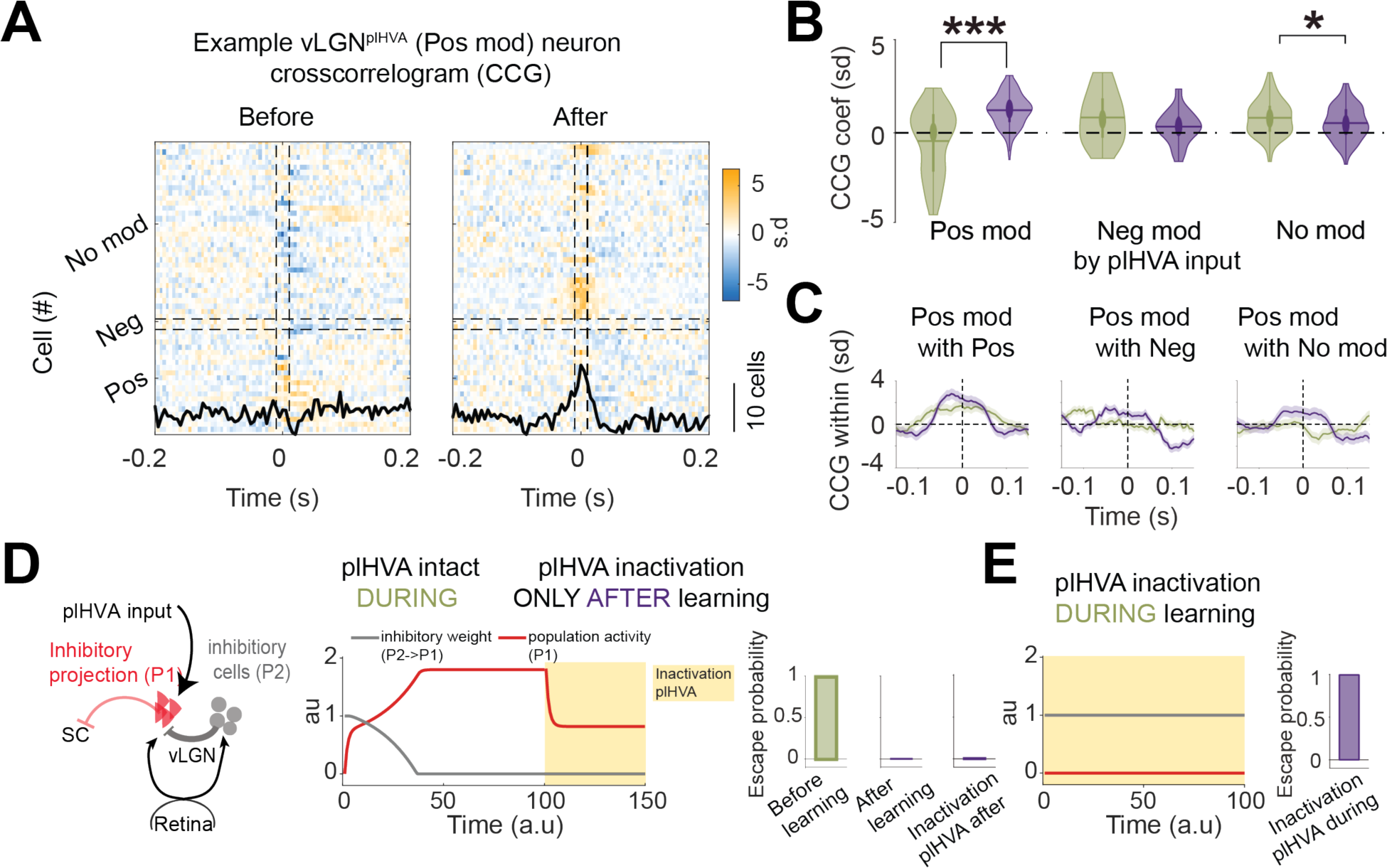
Inhibitory influence on vLGN^plHVA^ neurons may be decreased during learning. **(A)** Cross-correlogram (CCG) of temporal spiking relationships before and after learning between an example vLGN^plHVA^ neuron and all other simultaneously recorded units (each row shows a CCG, the spiking variation of the example neuron conditioned on the spiking of one other neuron at time 0. The black line shows the average of all CCGs in the plot. **(B)** Average CCG spiking variation around lag 0 (−1 ms to +4 ms) and distribution for all vLGN neurons positively, negatively or not modulated during plHVA optogenetic activation with the rest of the vLGN population before (green) and after (purple) learning. (before vs after learning, pos. mod.: P < 0.001; neg. mod: P = 0.785; no mod.: P = 0.011; KW test; n = 121; 29; 249 from 5 mice). **(C)** Average spike cross-correlogram for all vLGN neurons positively modulated by plHVA stimulation with other neurons from the three groups before (green) and after (red) learning. **(D)** Left, Schematic of the network model. Middle, population activity of vLGN^plHVA^ neurons (P1 in the model) and inhibitory weight change onto this population (from P2 onto P1) across learning (time 0 corresponds to steady-state before learning, and time 100 to after learning. Escape probability in the model from an average of 100 trials before and after learning. **(E)** Same as D but with removal of plHVA input during learning.

To explore if plastic changes in vLGN circuits could, in principle, arise from learning to suppress escape responses, we built a simple computational model simulating mean-field activity of two inhibitory neuronal populations both receiving sensory input from the retina (Fig. 4D). Population 1 (P1) consisted of neurons with long-range projections to SC that received input from plHVAs. Since previous work showed that high activity in SC-projecting vLGN neurons suppresses escape behavior, while low activity promotes fear responses (*11*, *34*) we equated low activity of P1 during presentation of looming stimuli with a behavioral escape response of the model and high activity of P1 with a suppression of looming-evoked escapes. In the model, this population received inhibitory connections from the second neuronal population (P2) that was not innervated by plHVAs, with plastic inhibitory weights. In steady-state, the inhibitory weights from P2 onto P1 were high, P1 activity was accordingly low, and the model exhibited escapes in response to looming stimuli. During subsequent training, the looming stimulus was paired with plHVA activation, which eventually resulted in increased activity of P1. This increase was accompanied by a gradual decrease in the effective inhibitory current from P2 to P1 and led to a cessation of escape behavior (Fig. 4D). Removing plHVA input during training prevented learning in the model, while removing plHVA input after learning had no effect on the learnt suppression of escape (Fig. 4E), recapitulating our experimental results.

The model exhibited depression of inhibitory synapses onto vLGN neurons that receive input from plHVAs during learning. Such a form of synaptic plasticity, called long-term depression of inhibition (iLTD), has been previously described in multiple brain areas *in vitro* and is dependent on endocannabinoid (eCB) signaling (*37–41*). Heterosynaptic iLTD can be triggered by activation of group I metabotropic glutamate receptors (mGluR1 or mGluR5) in postsynaptic neurons (*39*, *40*, *42*). This causes release of eCBs which act as retrograde messengers, activating eCB receptors (CB1R) on nearby presynaptic inhibitory terminals which can induce a long-lasting reduction of presynaptic GABA release probability (*37–41*). Allen Institute gene expression data showed that eCB receptor CB1R and mGluR5 are highly expressed in vLGN (Fig. S7A-B) (*43*). We therefore investigated if eCB-dependent iLTD in vLGN could mediate learned suppression of escape.

We first tested if learning to suppress fear responses was dependent on activation of mGluR5, by infusing mGluR5 antagonist MPEP in vLGN (Fig. 5A). Blocking mGluR5 receptors specifically in vLGN compromised learning: mice showed higher escape probabilities to looming stimuli after the learning protocol compared to vehicle-injected littermates (Fig. 5B). Blocking mGluR5 receptors in the hippocampus (dorsal of vLGN) instead had no effect on learning (Fig. 5B). Next, we examined if learning was mediated by eCB signaling. We infused a cocktail of eCBs synthesis inhibitors, LEI401+DO34, in vLGN (Fig. 5A). This intervention prevented animals from learning, as they still showed a high probability to escape from looming stimuli after the learning protocol (Fig. 5B). The same effect could be achieved by infusing a CB1 receptor antagonist (rimonabant) in vLGN (Fig. 5B), corroborating that eCB signaling is crucial for learning to suppress escape responses.

**Figure 5.**
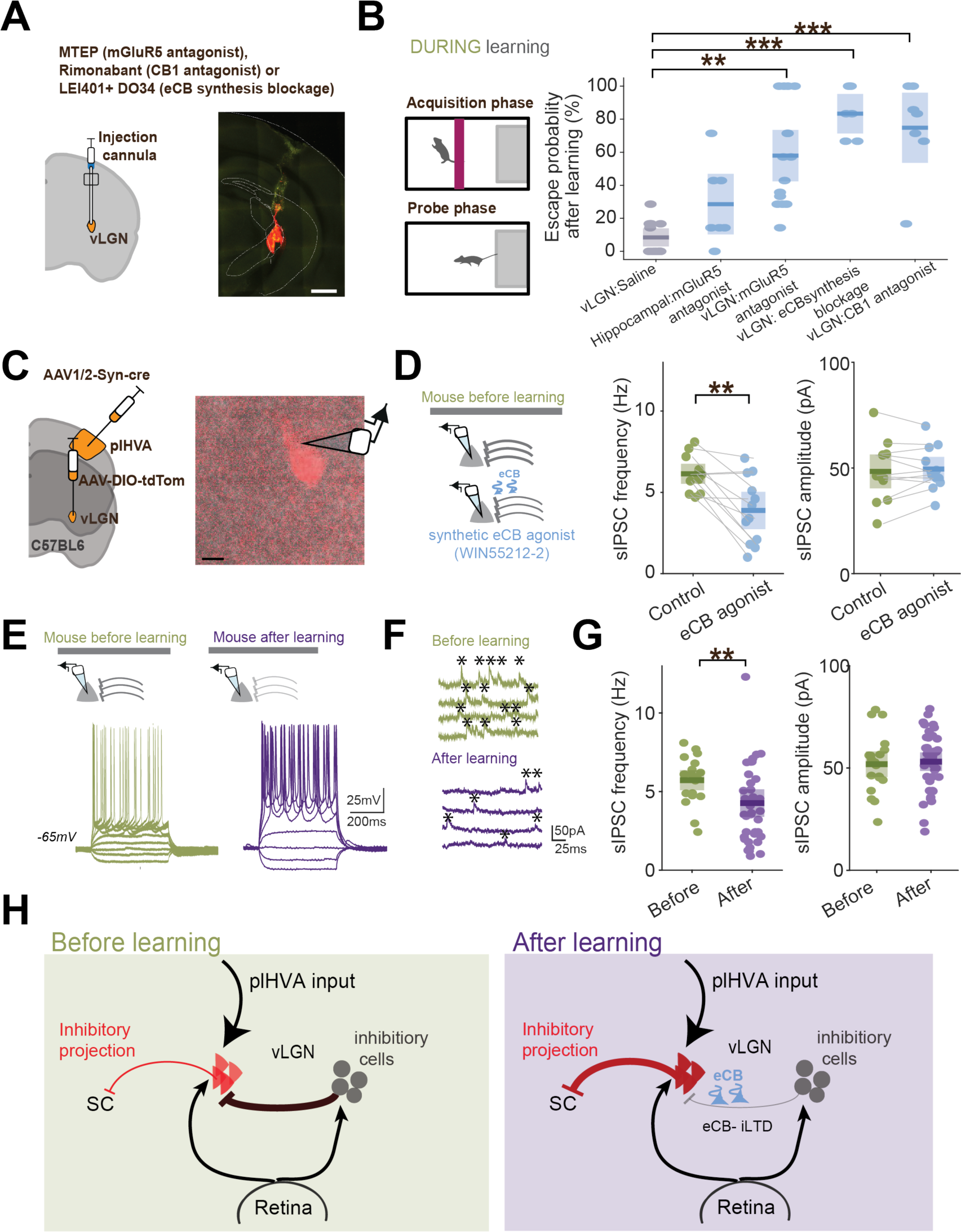
eCB-mediated long-lasting disinhibition as a mechanism for learning to suppress fear responses. **(A)** Left, Schematic of injection cannula implant in the vLGN. Right, Coronal section with an example injection. Scale bar 1 mm. **(B)** Left, Schematic of the task design. Right: Boxplot (with median and IQR) of post-learning escape probabilities for different reagents injected before the learning protocol; injection of saline in vLGN (13 mice), mGluR5 antagonist MTEP injection in hippocampus (P = 0.740, 7 mice), MTEP injection in vLGN (P = 0.001, 15 mice), eCB synthesis blockers LEI401 and DO34 injection in vLGN (P = 0.0004, 6 mice), and CB1 receptor antagonist Rimonabant injection in vLGN (P=0.0008, 7 mice). KW test followed by Tukey HSD for all conditions. Dots indicate individual animals **(C)** Left, experimental approach to visualize vLGN neurons receiving input from plHVAs (vLGN^plHVA^ neurons). Right, example of a recorded vLGN^plHVA^ neuron under the microscope. Scale bar 30 µm. **(D)** Left, Schematic of *in-vitro* whole-cell recordings from identified vLGN^plHVA^ neurons with bath application of eCB agonist WIN55212-2. Right, boxplots of frequency and amplitude of sIPSCs recorded with and without bath application of the eCB agonist (Freq: P = 0.003; Amp: P = 0.603; paired t-test; n = 12 cells from 4 mice). Dots indicate individual neurons. **(E)** Top, schematic of whole-cell recordings from identified vLGN^plHVA^ neurons in brain slices from mice before and after learning. Bottom, Membrane potential traces of example neurons in response to current injections of different amplitudes. **(F)** Example traces of spontaneous inhibitory postsynaptic currents (sIPSCs) under voltage-clamp in vLGN^plHVA^ neurons before (green) or after learning (purple). (G) Boxplots of frequency and amplitude of sIPSCs recorded from mice before and after learning (Freq P = 0.002; Amp P = 0.827 KW test; n = 20 cells from 4 mice before learning, 39 cells from 5 mice after learning). Dots indicate individual neurons. (H) Schematic of the mechanism underlying learnt suppression of escape responses: eCB-mediated inhibitory-LTD of presynaptic inhibition onto vLGN^plHVA^ neurons, likely induced via depolarization induced by direct visual looming stimulus input combined with input from plHVAs. Statistical significance is indicated as **P < 0.001 *** P < 0.001.

While our results so far indicate that eCB-dependent plasticity in vLGN underlies the learning process, it still remains open if and how synaptic connectivity in vLGN is altered by learning. We thus performed whole cell recordings in vLGN, specifically from vLGN neurons receiving input from plHVAs (vLGN^plHVA^ neurons, Fig. 5C-G). Bath application of eCB decreased the frequency, but not the amplitude of spontaneous inhibitory postsynaptic currents (sIPSCs) recorded from vLGN^plHVA^ neurons of naïve mice (Figure 5D), demonstrating eCB-induced suppression of inhibitory input onto these neurons. To identify learning-induced changes in vLGN circuits, we next performed whole-cell recordings of vLGN^plHVA^ neurons in mice that had undergone our learning protocol and learned to suppress escapes (Figure 5E). The excitability of vLGN^plHVA^ neurons was increased after learning (Fig. S7). However, more strikingly, we found that the frequency of sIPSCs in vLGN^lHVA^ neurons was significantly decreased after learning (Figure 5E-G), while their amplitude remained unchanged (Figure 5G). This indicates that release probability of GABA from presynaptic inhibitory terminals is decreased during learning, providing further support for eCB-dependent iLTD onto vLGN^plHVA^ neurons (*42*, *44*) (Figure 5H). Together, these results show that vLGN neurons receiving input from plHVAs increase their responses to looming stimuli over learning through a decrease in their inhibitory input via eCB-dependent plasticity (Figure 5H).

## Discussion

In this study, we uncovered a subcortical synaptic plasticity mechanism for learning to suppress instinctive defensive behavior instructed by a visual cortical area. Our findings highlight the critical role of the neocortex, specifically posterior lateral higher visual areas (plHVAs), in modulating instinctive fear responses based on experience. This is consistent with previous work showing that top-down projections from sensory cortex can influence instinctive behaviors and reflexes (*12*, *13*, *22*, *23*), and suggests that one evolutionary advantageous role of neocortical circuits could be to enable more flexible and adaptive behavior through the regulation of brainstem-driven instincts (*6–9*).

We find that higher-order visual cortex is essential for learning to suppress instinctive defensive reactions to visual threats. Surprisingly, visual cortex did not prove necessary for executing and sustaining the adaptive behavior once it was learned. This challenges traditional models which attribute learning and behavioral flexibility mainly to plasticity in telencephalic brain regions (such as neocortex, hippocampus, amygdala or striatum). While visual cortex activity may also change over learning, our results show that plasticity in these cortical circuits does not underlie the behavioral changes after learning. Instead, visual cortex activity crucially contributes to inducing experience-dependent plasticity down-stream, namely in the ventrolateral geniculate nucleus (vLGN). vLGN neurons driven by plHVAs increase their responses to the visual threat stimulus over learning, and such increased activity in vLGN circuits can abolish fear responses through their inhibitory influence on downstream areas that mediate escapes from visual threat such as the superior colliculus (*11*, *33*, *34*). The vLGN, and perhaps caudal prethalamic areas more generally (*33*, *35*), can thus link cognitive, neocortical processes with ‘hard-wired’ brainstem-mediated behaviors, providing a plastic inhibitory control pathway for experience-dependent adaptive behavior.

We find that learning to suppress fear responses relies on an endocannabinoid (eCB)-mediated form of inhibitory long-term synaptic plasticity, known as iLTD. This mechanism acts on inhibitory synapses onto vLGN neurons activated by plHVAs, decreasing presynaptic release probability. eCB-dependent iLTD has been demonstrated in multiple brain areas as an heterosynaptic *in vitro* plasticity mechanism induced through activation of glutamatergic mGluR5 receptors via repeated electrical stimulation in brain slices (*38–41*). We show that learning to suppress fear responses *in vivo* depends on mGluR5 receptor activation, endocannabinoid release and endocannabinoid receptor CB1 activation specifically in vLGN. Moreover, vLGN neurons receiving input from plHVAs, which are necessary for learning, are susceptible to eCB-dependent suppression of inhibition and show decreased inhibitory input after animals have learnt to suppress escape responses. Our study thus provides the first direct evidence of this plasticity mechanism - eCB-mediated decrease of inhibitory input - occurring in vivo to mediate learning. While the source of the plastic inhibitory input onto plHVA-driven vLGN neurons remains to be identified, it likely stems at least partly from local inhibitory interneurons: our cross-correlogram analysis suggests that plHVA-driven vLGN neurons are inhibited by other vLGN neurons before learning, but not after learning.

Consistent with our findings, eCBs have long been implicated in the regulation of fear and anxiety and are necessary for extinction of fear conditioning (*45*, *46*). Interestingly, both the suppression of instinctive fear responses and learnt fear extinction involve learned attenuation of defensive behaviors in response to a stimulus that no longer predicts danger (*47*). However, these two forms of learning likely engage distinct neural circuits and may differ in their specificity and time course (*46*, *48*). The plasticity mechanism described here is likely part of a larger network for regulating defensive behavior, including processes in downstream areas SC and PAG, as well as complementary top-down pathways through the basal ganglia, hypothalamus and amygdala (*20*, *21*, *48–53*).

The ability to suppress instinctive fear responses when threat expectations are violated is an ethologically crucial form of behavioral adaptation, whose absence could lead to inappropriate or excessive fear responses (*53*). Such maladaptive fear processing is a hallmark of fear and anxiety disorders and post-traumatic stress disorder (PTSD) (*52*, *54*). Dysfunction of pathways through vLGN (also called pregeniculate nucleus in primates (*55*)) or impairments in eCB-dependent plasticity could thus contribute to these disorders. Conversely, targeting these pathways or enhancing eCB-dependent plasticity within these circuits may facilitate suppression of maladaptive fear responses, suggesting novel therapeutic strategies for fear-related disorders.

In summary, we describe - from neural pathway to synaptic mechanism - a novel control mechanism that can overwrite instinctive responses when those are no longer advantageous. This adaptive behavior shaped by experience is mediated by endocannabinoid-dependent inhibitory plasticity in the diencephalon, instructed by visual cortex.

## Acknowledgments

We thank Michael Lohse, Alex Fratzl, and Manuel Valero for their feedback on the manuscript, Mrsic-Flogel and Hofer Lab members for helpful discussions, and Alex Fratzl for initial discussions, conceptualization and help with experimental setups and analysis. We thank Stephen C Lenzi and Troy W Margrie for help with the behavioural paradigm. We thank Michelle Li for animal husbandry and genotyping; Neurogears (Andre Almeida, Joao João Frazão and Gonçalo Lopes) for help building the behavioral setup; Alice M Koltchev and Lucie Daveau for help with animal handling; Rob Campbell for help with serial two-photon imaging; Piotr Nowak from the Sainsbury Wellcome Centre viral core for providing viruses. SWC FabLabs for technical support. Ana Covelo for advice on endocannabinoid experiments.

## Author contributions

Conceptualization: SM, SBH

Methodology: SM, SBH, NV

Investigation: SM, PB, NV

Experimental setup: SM, PB

Computational model conceptualization: CC, SM, SBH

Computational model investigation: CC

Funding acquisition: SM, SBH

Writing – original draft: SM, SBH

Writing – review & editing: SM, SBH, PB, NV, CC

## Data and materials availability

The data and code that support the findings of this study are available from the corresponding authors upon request.

## MATERIALS and METHODS

### Mice and ethics

All experimental procedures were performed in accordance with UK Home Office regulations (Animal Welfare Act of 2006), under project license PPL PD867676F & PP4459082, following local ethical approval by the Sainsbury Wellcome Centre Animal Welfare Ethical Review Body. Reporting has followed ARRIVE guidelines. Mice were housed in individually ventilated cages (IVC) under controlled climate (20-24°C; 45-65% humidity) in a 12:12 light:dark cycle with ad libitum access to laboratory food pellets and water. Wild-type C57BL/6J (Charles River), VGAT-IRES-Cre (Stock #028862, Jackson), VGAT-ChR2-EYFP (Stock #014548, Jackson), Ai14D (Stock # 007914 for cre-dependent tdTomato expression) and Rbp4-KL100 BAC-cre driver line (*56*) (MMRRC no. 031125-UCD; Selective labeling of layer 5 neurons) mice from either sex were used and were a minimum of 10 weeks old at the start of the experiments.

### Surgical procedures

#### Viral vector injection and fiber implantation

Prior to surgery, mice were subcutaneously injected with meloxicam (Metacam; 1 mg/kg). Mice were anesthetized with isoflurane (5% induction, 1.2–1.8% maintenance) in oxygen (0.9 L per min flow rate). Body temperature was maintained at 36.5°C using a controlled heating pad. The eyes were protected from light by aluminum foil and from drying by Xailin lubricating eye ointment. The mice were head fixed with ear bars on a stereotaxic frame (Kopf, model 940) and the scalp was cut along the midline to expose the skull. Small craniotomies (0.4-0.5 mm diameter) were made with a dental drill and AAVs were injected into the target brain regions using a pulled glass pipette (∼ 20 µm inner diameter at the tip; volume for each area indicated below) and a programmable nanoliter volume injector (Nanoject III, Drummond Scientific). AAVs were injected stereotaxically into vLGN (75nL per side, bregma: mediolateral: +/-2.5mm, anteroposterior: −2.3mm, dorsoventral: −3.5mm), plHVAs (120nL per side, midsaggital*: +/-3.75; transverse sinus*: +1.40; dorsoventral: −0.60; *Coordinates from midsaggital suture and transvere sinus). Ten minutes after the injection, the glass pipette was retracted and the incision was either sutured, or optic fibers (200 µm diameter, Doric Lenses) were implanted and fixed in place with light cured composite (RelyX Unicem 2 Automix from 3M). The optic fibres were implanted bilaterally above either vLGN (from bregma: mediolateral: +/-3.5mm, anteroposterior: −2.3mm, dorsoventral: −2.1mm, in a 20° lateral to medial angle to prevent penetrating the optic tract), plHVA (3.75 mm lateral to midline; 1.40 mm anterior to transverse sinus; 0.20 mm below dura), SC (mediolateral 0 mm, anteroposterior: −4.1 mm, dorsoventral: −1.7 mm, in a 30° posterior to anterior angle) or V1 (mediolateral 2.5 to 3 mm, anteroposterior: −3.3 to −3.5 mm, dorsoventral: −0.7 mm). The tip of the optic fibres were positioned 150 μm above target brain regions. After recovery, the animals were returned to their home cage for eighteen to twenty-nine days before experiments.

For pharmacological inactivation experiments, cannulae were bilaterally implanted over vLGN (3.06 mm lateral and 2.31 mm posterior to bregma, 3.01 mm below skull surface at a 10° lateral to medial angle) or plHVA (3.75 mm lateral to midline; 1.40 mm anterior to transverse sinus; 0.20 mm below dura). During surgery, each guide cannula was first inserted with an internal cannula with 1 mm projection, and dorsoventral coordinates were relative to the tip of the internal cannula. The internal cannulae were replaced with dummy cannulae with no projection to keep guide cannulae clear until intracranial injections were administered.

#### Viral constructs

We used AAV-EF1a-DIO-hChR2-EYFP (4.0 × 1012 vg/mL, 1:2 dilution, UNC vector core) for optogenetic activation of axon terminals; AAV1-hSyn1-SIO-stGtACR2-FusionRed (stGtACR2, Addgene: 105677-AAV1, 2 × 1013 vg × ml−1, diluted to 2 × 1012 vg × ml−1 in saline) (*57*) AAV2-EF1a-DIO-eNpHR3.0-EYFP-WPRE-pA (4.0 × 1012 vg/mL, 1:2 dilution, UNC vector core using Addgene plasmid #26972, a gift from Karl Deisseroth) for optogenetic silencing; AAV1-hSyn-DIO-hM4D(Gi)-mCherry (hM4Di, Addgene: 44362-AAV1, 1.8 × 1013 vg × ml−1, diluted to 9 × 1012 vg × ml−1 in saline) was used for chemogenetic inhibition; AAV1-hEF1a-mCherry (5.7 × 1012 vg/mL, 1:2 to 1:3 dilution, Zurich vector core) AAV-EF1a-FLEX-EGFP (GFP, 1 × 1014 vg ml−1, diluted to 2 × 1013 vg × ml−1 in saline) for control experiments for stimulation; hSyn-Flp-WPRE for data in Fig. S3 (SWC Vector Core, 2.5 × 1013 vg/mL). AAV2/1-hSyn-Cre-WPRE (1014 vg/ml). AAV2/5-EF1a-DIO-taCasp3-T2A-TEVp (1014 vg/ml. 1:3 dilution) for ablation.

#### Chronic implantation of silicon probes

All animals (28–35 g, 2–5 months old) were implanted with 64-site silicon probes (Cambridge NeuroTech) above the vLGN (mediolateral: +2.5mm, anteroposterior: −2.3mm, dorsoventral: −2.9 mm). Ground and reference wires were implanted in the skull above the cerebellum, and a grounded copper mesh hat was constructed, shielding the probes. Alternatively, a microdrive with the probe and ferrules were cemented to the skull protecting the area instead of using the copper mesh. Probes were mounted on microdrives that were advanced to the final recording vLGN area over the course of 5–8 d after surgery. For most of the experiments, a 200-μm optic fiber was attached to one of the shanks of the silicon probe to stimulate or inhibit the terminals from plHVA. A backup optic fiber was also implanted over plHVA. The back end of the fiber was coupled to a laser patch cord and stimulated as described for optogenetic manipulations. Animals were allowed to recover for at least 1 week before testing.

### Behavioral protocols

#### Escape behavior assay

Experiments assessing escape behavior in response to looming stimuli were performed in a custom-made transparent acrylic arena (80 cm × 26 cm × 40 cm (l × w × h)) with a red-tinted acrylic shelter (14 cm × 14 cm × 14 cm (l × w × h)) placed at one end (safe zone) and overhead looming stimuli presented at the other end (threat zone). The arena was placed in a large light-proofed and sound-attenuating box with a near-IR GigE camera (acA1300-60 gmNIR, Basler; with a frame rate of 50 fps) fixed on the ceiling to video record mouse behavior. To display looming stimuli, a projector (HF85JA, LG) was mounted in the box and back-projected onto a suspended horizontal screen (60 cm above arena floor, 100 cm × 80 cm; ‘100 micron drafting film’, Elmstock). The screen was kept at a constant background luminance level of 9 cd × m−2. The arena was illuminated by a strip of infrared LEDs to ease tracking of the mice. Video recording, looming stimuli and optogenetic laser stimulations were triggered through bonsai visual programming language (https://bonsai-rx.org/ and (*58*)). Mice were placed in the arena 10 minutes before the test to habituate to the new environment. Stimulation was triggered automatically when the mouse (tracked by bonsai) reached the threat zone. The stimulus was only triggered when the mouse was facing and walking toward the threat zone, with an inter-stimulus interval of > 90 seconds. The probability of stimuli being presented upon entering the threat zone was ∼50%. Each visual stimulation consisted of three consecutive expandinging black spots in a 3-second period. Each spot expanded from 1° to 20.8° in 330ms and was held at a constant size for 200ms. Optogenetic stimulation in laser trials started 0.2 seconds before the visual stimulus. Position and running speed of the center of the mouse were processed (after extraction with DeepLabCut, described below) using a custom-made MATLAB script. A successful escape was defined as a return to the shelter with an average running speed higher than 0.4 cm/s (varying this threshold to 0.3 or 0.5 cm/s did not change the significance of the results (*11*)) in the three seconds after the stimulus onset and reaching the shelter within five seconds of stimulus onset (two extra seconds for electrophysiological recordings). To qualify as escape, the mouse needed to be at least 10 cm away from the shelter, to be facing toward the threat zone (head direction at an angle less than 60 degrees from the front), and to reach the shelter in less than 2.5 s after escape onset. Escape onset was defined as the last time point before the onset of body rotation leading to an escape, defined as an angular velocity > 100 deg/s.

For electrophysiology experiments, looming stimuli of the highest contrast (100%) were presented until the mouse stopped escaping, on average after 23 +/- 14 stimulus presentations (mean +/- sd). After 3 consecutive high contrast looms without escapes, the animal was considered to have learned to suppress escape responses. Once this threshold was reached the animal was presented with 15-20 more looms.

#### Escape behavior assay with wall partition

An optional red Perspex partition (wall) could be inserted to isolate the “threat end” of the arena, 28 cm from the threat-end wall. For these experiments mice were introduced to the threat zone of the behavioral arena with the wall preventing the mouse from seeing or reaching the shelter. Mice were habituated to the arena with the wall in place for 10 min before stimulus onset. During an acquisition phase, looming stimuli were presented with an inter-stimulus-interval of > 90 seconds at increasing contrast levels, in sets of three stimulus presentations, starting with three stimuli at 20% contrast, and repeated for six further increasing contrast levels (40%, 60%, 80%, 90%, 100%). During the test phase mice faced 6-10 repetitions of 100% contrast looming stimuli. Mice were considered habituated if their response remained consistent for the last 3 of 6 trials. If not, testing continued up to 10 trials to confirm the outcome.

### Extraction of behavioral variables

Raw videos of behavioral experiments and corresponding input channels (laser, visual stimuli and camera trigger) were acquired using Bonsai software. The camera was triggered at 1Hz, with analogue input channels recorded at 50 samples/s. Timestamps of visual stimuli and laser onsets were extracted using a MATLAB script and associated with the corresponding video frame.

To extract behavior we used a deep neural network approach (DeepLabCut; (*59*)) as described previously (*11*). A network was trained with 100 manually annotated frames of an initial dataset, and then re-trained with 25 new frames each from two additional datasets from different mice. The resulting labeled videos were manually inspected for quality control. This network was used for all behavioral classifications, and the XY-position of body-center, head, nose, and ears within the arena were extracted. XY-positions were loaded and processed in MATLAB. The position of the animal was defined as the body-center position. The head-direction of the animal was computed as the vector between the nose and the head (0 degrees: mouse facing toward the “threat zone”; ± 180 degrees: mouse facing toward the shelter). Instantaneous speed was computed as the Euclidean distance between two positions at consecutive time points. Angular velocity was computed as the discrete derivative at two consecutive time points (taking into account circular boundary conditions). Prior to computing the speed and angular velocity, the position and head direction vectors were smoothed by a 100 ms running average filter to avoid amplifying noise when computing derivatives. The ears were tracked to increase the stability of the network with more co-dependent markers. For all experiments, data analysis was performed in custom-written routines in MATLAB (Mathworks).

### Optogenetic manipulation

Optogenetics Laser stimulation protocols were created and run through custom-made scripts in Bonsai and commanded to a Pulse Pal pulse generator (Open Ephys). For stimulation of ChR2 and stGtACR2 (*59*), a 473 nm laser (OBIS 473nm LX 75mW LASER SYSTEM, Coherent) was used, coupled to a 200 µm fiber patch cable through an achromatic fiber port (Thorlabs). For stimulation of eNpHR3.0 we used a 594 nm laser (OBIS 594nm LS 40mW LASER SYSTEM: FIBER PIGTAIL: FC, Coherent) instead. The laser frequency was 20 Hz (with 50% duty-cycle pulses) for depolarizing opsins (ChR2) and 0 Hz for hyperpolarizing opsins (stGtACR2 and eNpHR3.0). The peak laser power at the tip of the fiber was ∼2.5 mW. To control for unspecific effects on behavior due to laser stimulation, we performed laser stimulation of control mice expressing EGFP or tdTomato.

### Pharmacological inactivation

Mice were bilaterally implanted with guide and dummy cannulae (Plastics One) over vLGN or plHVA (see Surgical procedures) and given at least 1 week for recovery. On the day of the experiment, mice were handled and an internal cannulae projecting 0.2 mm below the guide cannulae were inserted and fixed on awake animals. Drug or vehicle were then infused at a rate of 150–200 nl per minute using a microinjection unit (Hamilton, 10-μl syringe) connected to the internal cannulae through tubing (Plastics One) and plastic disposable adapters (Plastics One). For inactivation of plHVAs, 0.5-0.6 μl Muscimol-BODIPY-TMR-X (0.5 mg ml−1, ThermoFisher) or vehicle was injected into plHVAs in each hemisphere. For vLGN infusions, 0.2-0.3 μl drug or vehicle was injected per vLGN hemisphere. MTEP (CAS 329205-68-7,1 mg); Rimonabant (National Institute on Drug Abuse, Rockville, MD 0.257 mg), and the cocktail of LEI401 (MedChemExpress (Monmouth Junction, NJ 0.5 mg) and DO34 (Aobious (Gloucester, MA) 0.5 mg) were infused into vLGN delivered in a vehicle composed of 1:1:18 of DMSO:Tween80:saline. For chemogenetic silencing experiments, water-soluble clozapine N-oxide dihydrochloride (CNO, Hellobio) was applied at a final concentration of 10 µM from a 100 mM stock solution. Vybrant CM-Dil Cell labeling solution (DiI; Thermo Fisher V22888) was added to the solution to confirm the site and extent of infusion upon tissue processing (described below). Mice were given 10 min to recover before starting the behavioral assay. Immediately upon completion of the behavioral assay, mice were anesthetized with pentobarbital (IP, 80 mg mg/kg) and transcardially perfused, as described below.

### Anatomical confirmation

After experiments were completed, mice were deeply anesthetized and transcardially perfused with cold 1M PBS followed by cold 4% paraformaldehyde. The brain was extracted and fixed for 24 h at 4°C in paraformaldehyde before being transferred to PBS at 4°C. The tissue was processed and imaged using 2-photon serial sectioning microscopy (BrainSaw, https://swc-advanced-microscopy.github.io/facility_webpage/). The microscope was controlled by ScanImage Basic (Vidrio Technologies, USA) using BakingTray, a custom software wrapper for setting up the imaging parameters (https://github.com/SainsburyWellcomeCentre/BakingTray, https://doi.org/10.5281/zenodo.3631609). Images were assembled using StitchIt (https://github.com/SainsburyWellcomeCentre/StitchIt, https://zenodo.org/badge/latestdoi/57851444)). Brains were registered to the Allen Common Coordinate Framework (*60*, *61*). Probe trajectories and locations of cannula injections and virus expression were reconstructed from images using publicly available custom code (http://github.com/petersaj/AP_histology & (*62*)).

### Extracellular electrophysiological recordings

An Intan RHD2000 system (Intan Technologies) was used to acquire electrophysiological data, which were digitized at 30kHz. The wideband signal was downsampled to 1250 Hz. A pulley system was developed to counteract the weight of the tether and headstage. In the open arena, optogenetic ChR2-stimulation was administered by 5 ms square pulses, which were repeated every 30 s for 1500 trials using PulsePal2 (Sanworks, v2). Square pulse stimulation for eNpHR3.0 was triggered by TTLs from using PulsePal2 controlled by Bonsai during the escape assay, starting 0.2 s before visual stimulus presentation and lasting until the end. To track the animal’s position, the setup controlled by Bonsai and described in the behavioral protocols was used, sampled at 30Hz. Position data was synchronized with neural data using TTLs that signaled the events triggered by Bonsai in the behavioral setup.

#### Unit clustering and neuron classification

Spike sorting was performed semi-automatically with KiloSort (https://github.com/cortex-lab/KiloSort (*63*)), using a pipeline KilosortWrapper (a wrapper for KiloSort, https://github.com/brendonw1/KilosortWrapper). Units were manually curated using the software Phy (https://github.com/kwikteam/phy) and plugins for Phy (https://github.com/petersenpeter/phy-plugins) (*64*). The following parameters were used for the Kilosort clustering: ops. Nfilt: 6 * numberChannels; ops.nt0: 64; ops.whitening: ‘full’; ops.nSkipCov: 1; ops.whiteningRange: 64; ops.criterionNoiseChannels: 0.00001; ops.Nrank: 3; ops. nfullpasses: 6; ops.maxFR: 20000; ops.fshigh: 300; ops.ntbuff: 64; ops.scaleproc: 200; ops.Th: [4 10 10]; ops.lam: [5 20 20]; ops.nannealpasses: 4; ops.momentum: 1./[20 800]; ops.shuffle_clusters: 1. Units were only included in any of the following analyses if they had a minimum firing rate of 0.5 Hz and were recorded from channels within vLGN as determined by anatomical confirmation.

We extracted binned z-scored firing rates (2ms bin size) from −2 to 5 s around looming stimulus onset, smoothed with a causal half Gaussian filter (standard deviation of 15 ms), to determine looming stimulus responses of vLGN units. Responses to all looming stimuli before (before learning) and after mice did not escape to three consecutive looming stimuli (after learning) were averaged. Bootstrap confidence intervals (1500 repetitions) were determined across trials and firing rate responses to looming stimuli were deemed significant if test–retest covariance value was above the 95th percentile of the shuffled distribution. Neurons were included in the analysis if their firing rate 0-300 ms post stimulus onset was higher or lower than the baseline plus 2 standard deviations in at least 15 consecutive bins, before or after learning or both. Trials in which escape initiation occurred before 300 ms were excluded from the analysis (1.5 % of trials).

Cell metrics were defined as follows by (https://cellexplorer.org/ (*65*)).

#### Optogenetics analysis during unit recordings with ChR2

To test if units were responsive to optogenetic activation (*66*) light-triggered post-event histograms were computed, binned at 1 ms width. If the peak firing rate in a 1 - 70 ms window after the light delivery (average of the first and second maximum value) was more than 5 standard deviations above the mean of the baseline period (−100 - 0 ms before the light delivery), the unit was defined as optogenetically activated (positively modulated). Neurons that were on average significantly suppressed during the window of light stimulation, with a p-value below 0.01 (ANOVA), were defined as negatively modulated.

#### CCG analysis

We implemented a cell-by-cell cross-correlation analysis to quantify temporal spiking relationships between all recorded cells in vLGN classified for optogenetic activation (see above). We excluded all looming stimulus presentations (5 s around each stimulus presentation) to minimize stimulus-driven correlations, and divided the recordings into two periods before and after learning. Average unit cross-correlograms (CCGs) were computed as the mean from all z-scored CCGs (5-ms binned, 0.6 s window), either with all other simultaneously recorded neurons or neurons of specific subgroups. The average CCG spiking variation was computed around lag 0 in a window from −1 ms to +4 ms.

### Computational Modeling

We simulate a mean-field model of our vLGN^plHVA^ population, 𝑃1, following the given dynamics

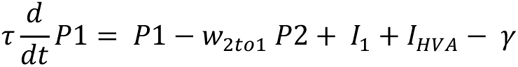

where 𝜏 = 2 [a.u.] is the time constant of the dynamics, 𝛾 = 0.2 𝑖𝑠 𝑎 𝑏𝑎𝑐𝑘𝑔𝑟𝑜𝑢𝑛𝑑 𝑐𝑢𝑟𝑟𝑒𝑛𝑡, 𝐼_1_ = 1 are the sensory currents into the P1 population, 𝐼*_HVA_* is the current from HVA, and −𝑤*_2to_*_1_ 𝑃2 is the current from a second inhibitory population P2 with 𝑃2 = 1. 𝐼*_HVA_* is 1 if HVA is active and 0 if it is inactive. P1 is restricted to be positive. The effective inhibitory currents are denoted by 𝑤*_2to_*_1_ from population 2 to 1. The effective mean-field weights are subject to the following plasticity rule

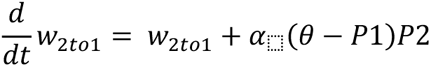

where 𝛼 = 0.05 is the plasticity learning rate and 𝜃 = 0.5 is a plasticity threshold. The weights are bounded, with a lower bound at 0 and an upper bound at 1.

We start by a warm-up simulation to bring the network to steady-state, where we initialize P1 = 0 and 𝑤*_2to_*_1_ =0.5 and simulate for 100 time steps. After the warm-up period and before learning, 𝑃1 is suppressed and 𝑤*_2to_*_1_ is high. We model the escape behavior as b = 2 − P1 with escape occurring at values lower than 0.5. Therefore before training, the model is escaping. During training, we assume 𝐼*_HVA_* = 1 and we simulate for another 100 time steps. During testing, we fix the weights, and we either set 𝐼*_HVA_*= 1 or 0 and simulate for an additional 100 time steps. For the condition where we have no HVA input during training, we set 𝐼*_HVA_* = 0 for training and keep all the other parameters the same for the 100-time step simulation.

### In vitro whole-cell recordings

#### Preparation of acute vLGN slices

Male and female mice of 6 to 8 weeks old were injected with AAV2/1-hSyn-Cre in plHVA and AAV-EF1a-FLEX-tdTomato in vLGN. After 3 to 4 weeks mice were perfused with ice-cold (4 °C) NMDG-HEPES solution prior to brain extraction following anaesthesia, to improve slice viability in adult mice (>1.5 months old). Brains were quickly removed and immediately immersed in ice-cold slicing solution NMDG-HEPES containing [in mM]: NMDG 92, KCl 2.5, NaH2PO4 1.2, NaHCO3 30, HEPES 20, glucose 25, thiourea 2, Na-ascorbate 5, Na-pyruvate 3, CaCl2·2H2O 0.5, and MgSO4·7H2O 10, with an osmolarity of 281-282 mOsm, and constantly bubbled with carbogen (95% O2 and 5% CO2) for a final pH of 7.3. Acute slices of 250 µm thickness were prepared containing the vLGN using a vibratome (Leica VT1200 S). Subsequently, slices were placed in the same NMDG-HEPES solution at 37 °C for 10 min. Slices were then transferred to a different recovery chamber and submerged in artificial cerebrospinal fluid (aCSF) solution containing (in mM): 125 NaCl, 2.5 KCl, 26 NaHCO3, 1 NaH2PO4, 10 glucose, 2 CaCl2, and 1 MgCl2, with an osmolarity of 293–298 mOsm and constantly bubbled with carbogen (95% O2 and 5% CO2) for a final pH of 7.3. Slices were further incubated at room temperature (19–23 °C) for at least 60 more minutes prior to electrophysiological recordings.

#### Whole-cell recordings

Pipettes were pulled from standard-walled filament-containing borosilicate glass capillaries (Harvard Apparatus, 1.5 mm outer diameter, 0.85 mm internal diameter) using a vertical micropipette puller (PC-10, Narishige) to a final resistance of 4–6 MΩ. Pipettes were backfilled with potassium methane sulfonate based solution containing (in mM): 130 KMeSO4, 10 KCl, 10 HEPES, 4 NaCl, 4 Mg-ATP, 0.5 Na2-GTP, 5 sodium phosphocreatine, 1 EGTA, with an osmolarity of 294 mOsm, filtered (0.22 µm, Millex) and adjusted to pH 7.4 with KOH.

Data was acquired using an Axon Instruments Multiclamp 700B amplifier and a National Instruments board. Recording pipettes were mounted on remote-controlled motorized micromanipulators (MicroStar, Scientifica). Cells were approached under visual guidance using laser-scanning Dodt contrast and two-photon imaging with a Scientifica MP-1000 multiphoton imaging microscope (in order to identify fluorescently labeled vLGN cells receiving plHVA input), a mode-locked Ti:sapphire laser (Vision-S, Coherent) and a Nikon 16x water-immersion objective (NA 0.8). Scanning and image acquisition were controlled by SciScan (Scientifica) and custom software written in LabVIEW (National Instruments). We used WinWCP 5.3.4 software for acquisition (developed by John Dempster, freely available at http://spider.science.strath.ac.uk/sipbs/software_ses.htm).

The resting membrane potential was determined in current clamp directly after establishing whole-cell configuration experiments were only continued if cells had a resting membrane potential < −45 mV. Cells were characterized by recording their membrane responses to current injections of increasing strengths (−40 to + 140 pA, increment: 20 pA, 500 ms duration). Voltage-clamp recordings were performed at a holding potential of 0 mV to record spontaneous inhibitory postsynaptic currents.

Pharmacology. To block glutamatergic synaptic transmission, a cocktail of antagonists was used, including D-(-)-2-amino-5-phosphonopentanoic acid (D-AP5; 50 µM) and 6-cyano-7-nitroquinoxaline-2,3-dione disodium (CNQX; 10 µM). In a subset of experiments, endocannabinoid agonist WIN 55,212-2 (5μM) (Tocris Bioscience) was bath-applied and after a resting period of 15 min, recordings were continued.

#### Analysis

Electrophysiological data were analyzed offline using custom-written functions in MATLAB after converting data from WinWCP. For whole-cell recordings, the resting membrane potential was calculated during current steps in a 120 ms baseline window in current clamp mode. The input resistance was calculated from the steady-state voltage measured in response to a hyperpolarising test pulse (−40 pA, 500 ms duration) at holding potential of −65 mV. The membrane time constant was calculated by fitting a single exponential to the decay of the test pulse (y = y0 + Ae(-(x-x0)/τ)). Rheobase was defined as the minimum amount of current required to evoke a single AP. Spontaneous inhibitory postsynaptic currents (sIPSCs) onto neurons were detected using a threshold algorithm, and their frequency and peak amplitude were analysed during 7 min. Statistical tests were performed on all cells pooled across animals.

### Statistics

All statistical tests were conducted using MATLAB R2021a, and the details of the tests used are described in the figure legends. All statistical analyses were performed with standard MATLAB functions. No specific analysis was used to estimate minimal population samples, but the number of animals, trials, and recorded cells were similar to or larger than those employed in previous work (*11*, *69*). Data collection was not performed blinded to the subject conditions. Data analysis was performed blinded to the scorer or did not require manual scoring. All subjects underwent the same number of conditions (unless stated otherwise) in a randomly assigned fashion. Unless otherwise noted, for all tests, non-parametric two-tailed Wilcoxon’s paired signed-rank test, Kruskal-Wallis, one-way analysis of variance and Friedman test were used. When parametric tests were used, including two-ways ANOVA, repeated-measures ANOVA the data satisfied the criteria for normality (Kolmogorov–Smirnov test) and equality of variance (Bartlett’s test for equal variance). For multiple comparisons, Tukey’s honesty post hoc test was employed and the corrected *P < 0.05, **P < 0.01, ***P < 0.001 are indicated. Results are displayed as Violin plots, representing median, 25th/75th percentiles, whiskers data range and the probability density of the data, unless indicated otherwise. Boxplots represent median, 25th/75th percentiles, whiskers data range. Dispersion represents ± IC95. The exact number of replications for each experiment is detailed in the text and figures.

