## Supplementary Figures and text for "Overwriting an instinct: visual cortex instructs learning to suppress fear responses"

**The PDF file includes:**

Supplementary Text
Figs. S1 to S7

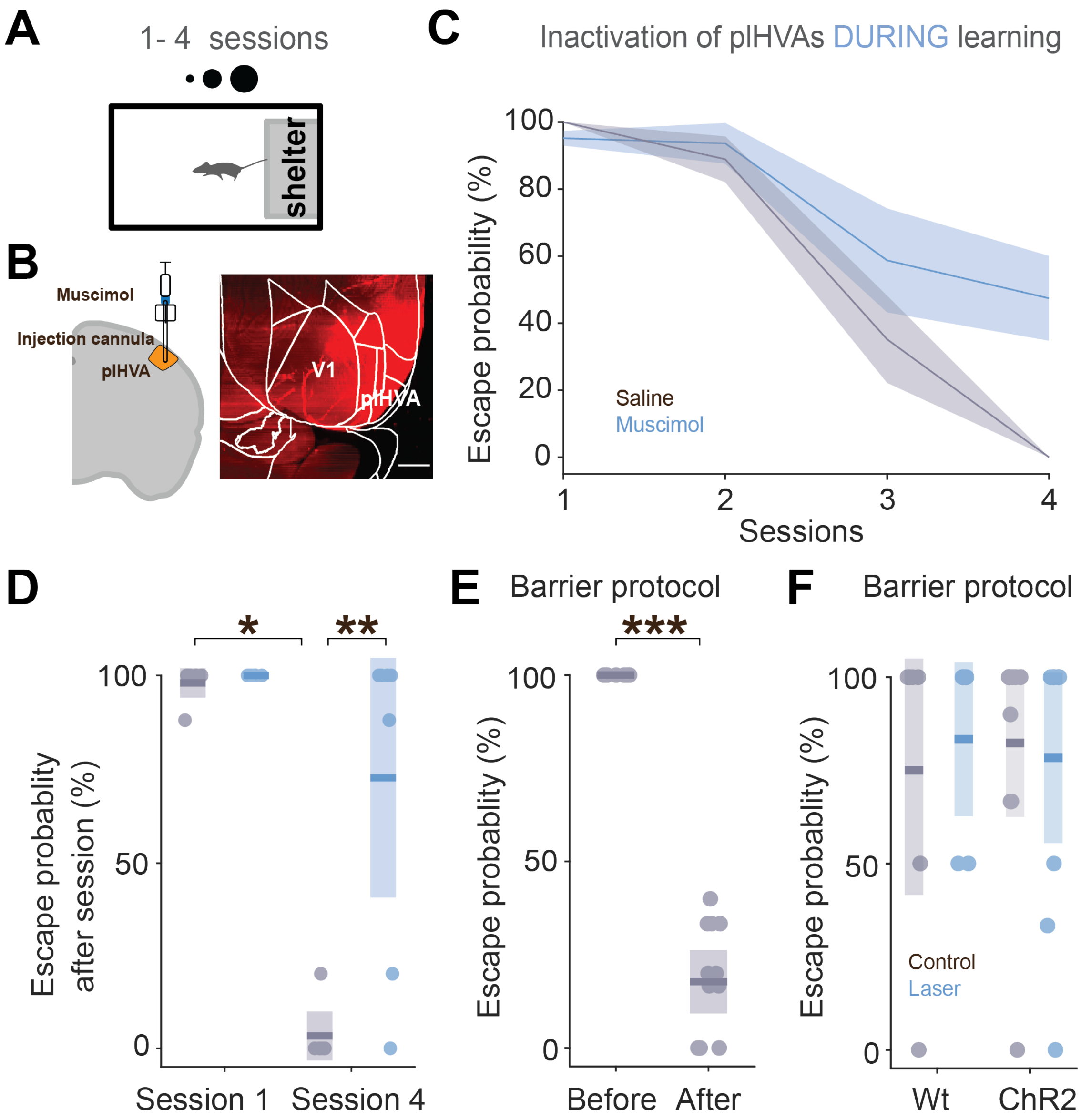


**Figure S1. plHVA activity is crucial for learning to suppress fear responses and does not influence instinctive responses, related to Figure 1. (A)** Variation of the training paradigm in which animals learn through repeated presentation of 100% contrast looming stimuli with access to the shelter in 1-hour sessions over days. **(B)** Schematic of the experimental approach for acute inhibition with Muscimol through an injection cannula targeting plHVAs. Scale bar 1mm. **(C)** Effects of Saline/Muscimol infusion in plHVAs on mean escape probability over a period of days, with shaded areas representing 95% confidence intervals. **(D)** Boxplots of escape probability after training on day 1 and day 4, with individual animals represented as pale dots. Day 1: $P = 0.9774$; day 4: $P = 0.002$, repeated-measures ANOVA, Saline: $P = 0.0152$; Muscimol: $P = 0.1923$; KW $n = 7$ mice. **(E)** Boxplot (showing median and IQR) of escape probability before and after the acquisition phase of the short learning protocol with barrier (see figure 1A). Pale dots represent individual animals. $P < 10^{-6}$, KW test; $n = 12$ mice. **(F)** Boxplots showing escape probability when plHVA are silenced during presentation of 100% contrast looming stimuli to naive animals before learning. All P-values > 0.998, KW test; n = 6 (WT) and 10 mice (VGAT: ChR2). Statistical significance is indicated as * P < 0.05; ***P < 0.001.

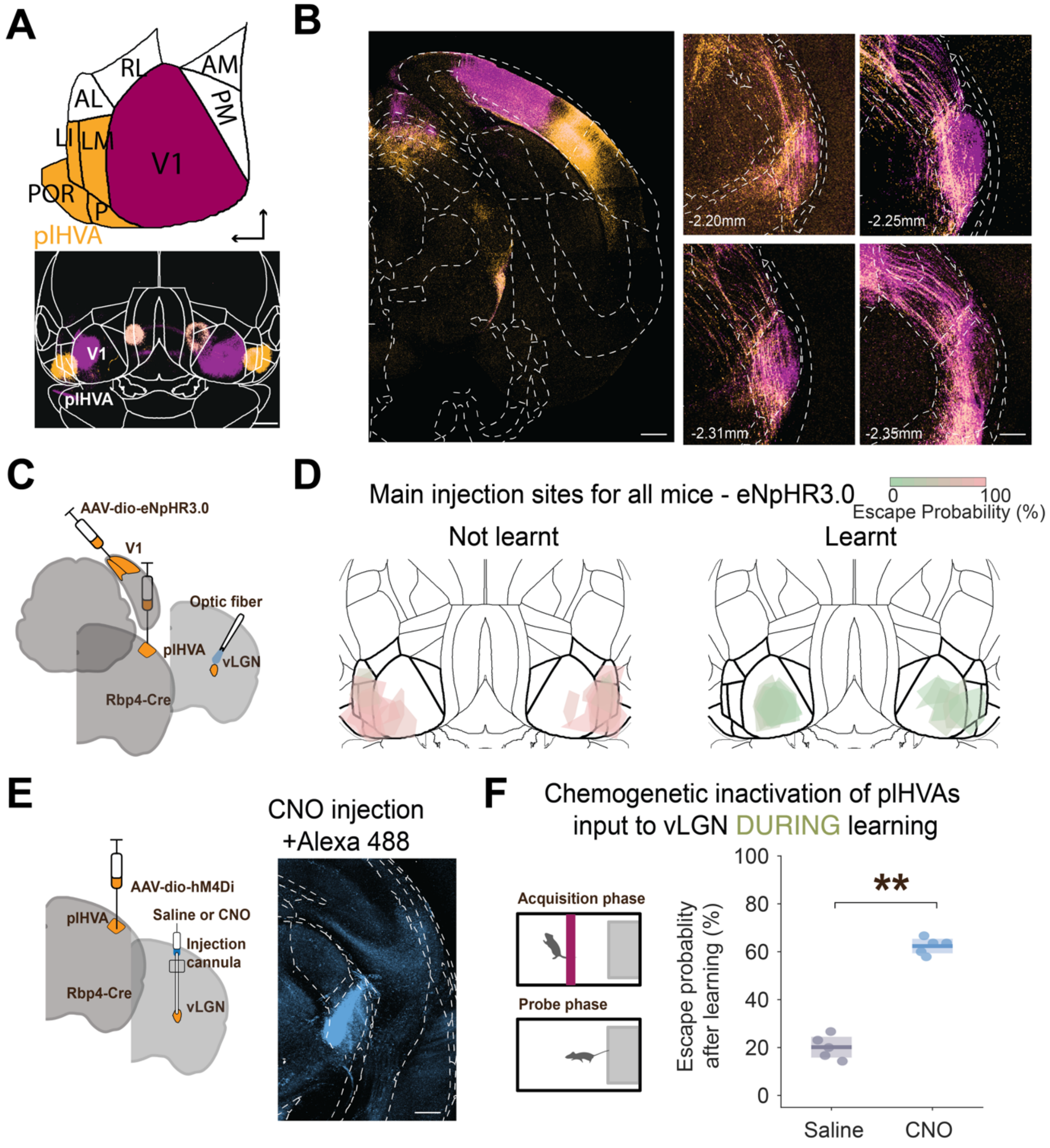


**Figure S2. Axons over vLGN from visual cortex, Related to Figure 1. (A)** Top: dorsal view of neocortical visual brain areas. Bottom, dorsal view of injection sites in V1 and plHVA with different tracers.

Scale bar 500 µm. V1, Primary visual cortex; PM, posteromedial; AM, anteromedial; RL, rostrolateral; AL, anterolateral; LI, laterointermediate; LM, lateromedial; POR, postrhinal; P, posterolateral. **(B)** Coronal view of cortical injection sites (left) and axon label patterns in vLGN and surrounding areas (right) resulting from injections in V1 (purple) and plHVAs (orange). Scale bar 1mm, 150 µm. Distance from bregma is indicated in the bottom left. **(C)** Experimental approach: eNpHR3.0 injections into visual cortical areas and optic fiber located above vLGN. **(D)** Dorsal views of neocortical areas with outline of injection sites for animals that learned to suppress escape responses after training (right), and mice that did not learn (left). **(E)** Left, experimental design: hM4Di was expressed in plHVAs, and CNO or saline injected into vLGN. Right, example image showing Alexa 488 fluorescence after CNO infusion in vLGN. Scale bar 150 µm **(F)** Left, Schematic of the task design. Right, boxplots (showing median and IQR) of post-learning escape probability, demonstrating the effects of injecting Saline or CNO into vLGN before the acquisition phase. $P = 0.008$, KW test; $n = 5$ and 5 mice. Statistical significance is indicated as ** $P < 0.01$. Individual animals are indicated by dots.

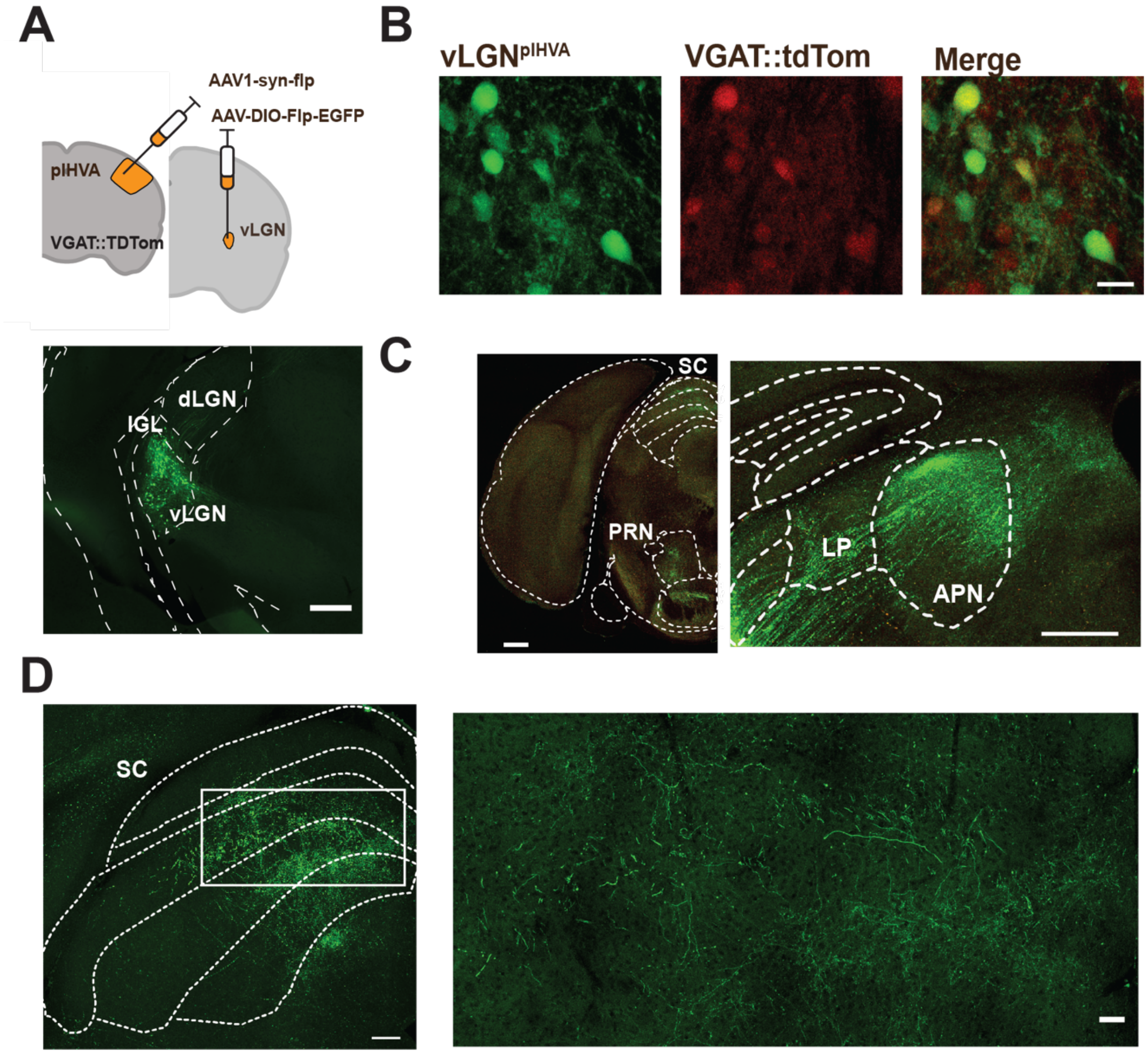


**Figure S3. vLGN$^{plHVA}$ cells are VGAT-positive and project to the SC. (A)** Experimental approach of anterograde transneuronal labeling. Bottom, example image of vLGN neurons receiving input from plHVAs (vLGN$^{plHVA}$ neurons). Scale bar 500 µm **(B)** Example images of EGFP expressed in vLGN$^{plHVA}$ and tdTomato expressed in vLGN VGAT+ neurons. Scale bar 20 µm. **(C,D)** Coronal views of axons of vLGN$^{plHVA}$ neurons in downstream areas. Scale bars 500 µm, 500 µm**,** 200 µm, 50 µm. vLGN: ventral lateral geniculate nucleus, IGL: Intergeniculate leaflet, dLGN: dorsal geniculate nucleus, PRN: pontine reticular nucleus, SC: superior colliculus, LP: lateral posterior nucleus, APN: Anterior pretectal nucleus.

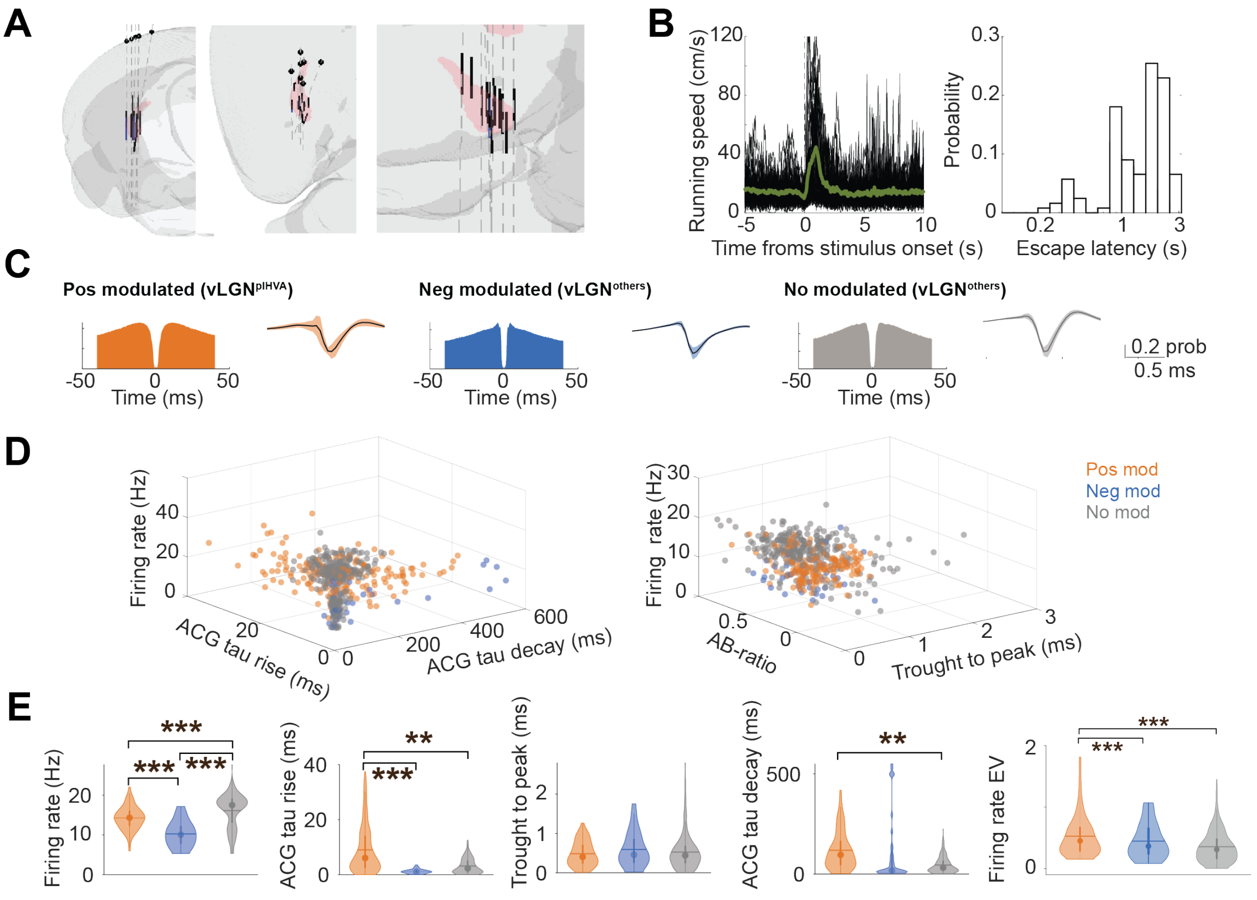


**Figure S4. Cell-type identity of vLGN cells. Related to Figure 3. (A)** Locations of silicon probes used in the experiment. **(B)** Left, escape running speed over trials before learning for one example mouse. Green trace shows the mean. Right, histogram of escape latencies for all animals and trials (122 trials from 9 mice). **(C)** Left, average firing auto-correlogram (ACG, left) and average spike waveform (right, mean ± IC95) for vLGN neurons positively (blue), negatively (red) and not modulated by plHVA activation (gray). **(D)** Parameters of waveforms and firing rates for each neuron plotted against each other. **(E)** Violin plots comparing different electrophysiological properties for the three groups of vLGN neurons, including firing rate across the recording session, autocorrelogram (ACG) tau rise, through-to-peak spike duration, ACG decay, and firing rate estimated variance (EV). ($P < 10-7$ for firing rate, KW test; $P < 0.0001$ and $P=0.0034$ for ACG tau rise, KW test; $P = 0.004$ for ACG tau decay, $P<0.001$ for firing rate std, KW test; n = 121; 29; 249 cells from 5 mice). ***$P < 0.001$, ** $P < 0.01$.

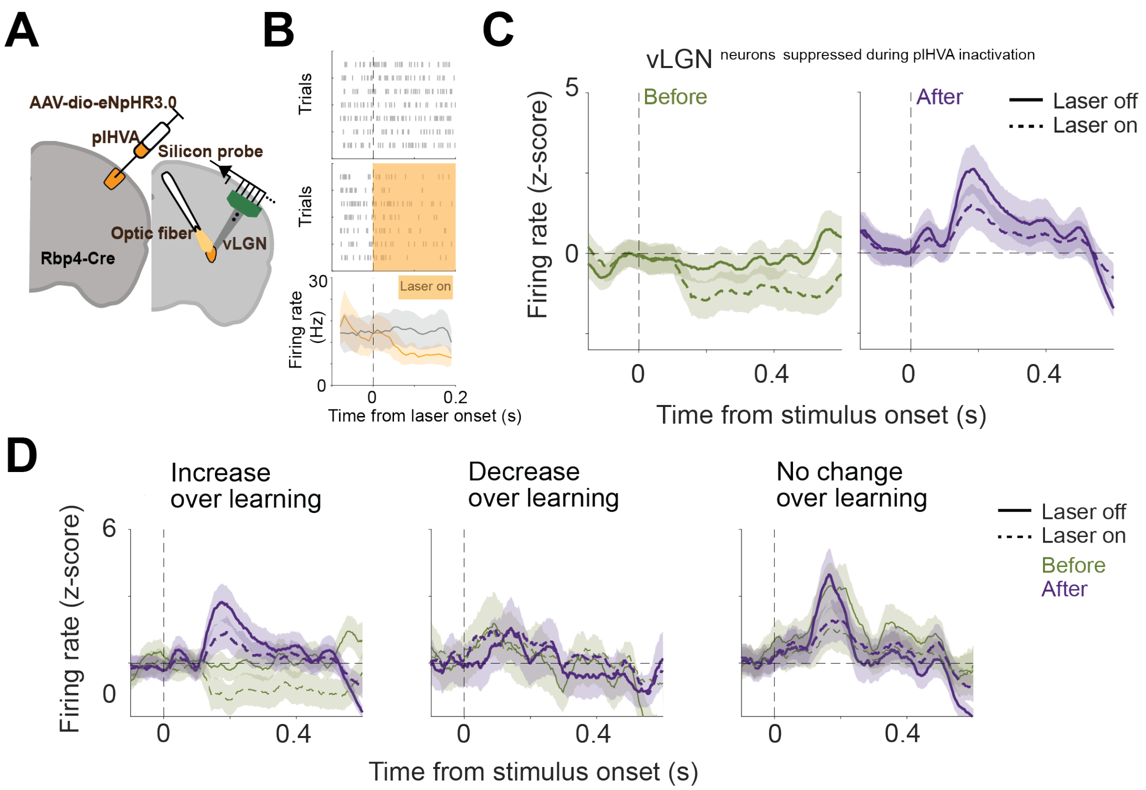


**Figure S5. Effect of plHVA silencing on vLGN responses to looming stimuli. (A)** Experimental design showing silicon probe electrophysiological recordings from vLGN cells combined with plHVA silencing. **(B)** Example neuronal response in trials without and with laser (yellow) for a neuron suppressed during plHVA inactivation. **(C)** Average PSTHs in response to looming stimuli with and without plHVA silencing before (left) and after (right) learning, for the vLGN neurons significantly suppressed during plHVA silencing (n = 116 from 4 mice). During laser stimulation trials, the laser was on from - 0.02 s before visual stimulus onset to + 3 s. **(D)** Average PSTHs in response to looming stimuli before and after learning with (dashed line) and without laser stimulation, for vLGN increasing, decreasing or not changing their looming responses over learning (n = 116; n = 85; n = 31 cells from 4 mice).

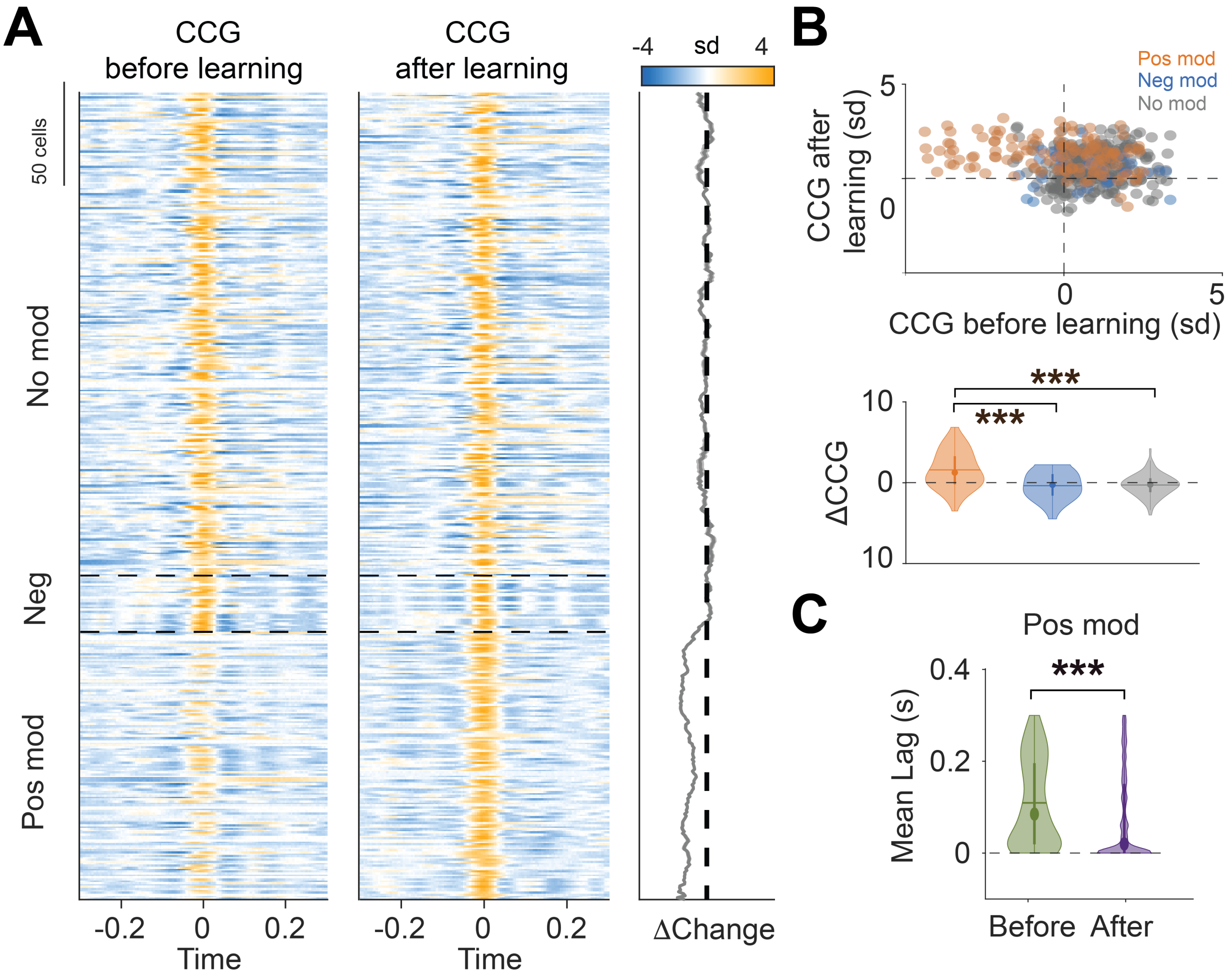


**Figure S6. Changes in spiking relationships between vLGN neurons over learning. (A)** Each row shows the average cross-correlogram (CCG) for an individual vLGN neuron with all other simultaneously recorded neurons before (left) and after learning (right). Neurons are ordered according to how they are modulated by plHVA activation. Right, difference in the CCG spiking variation around lag 0 (-1 - +4ms) before and after learning for each neuron. **(B)** Top, magnitude of average CCG response variation around 0 time lag (-1 - +4ms) after learning for each cell as a function of its CCG response variation before learning. Bottom, difference between the values above of average CCG response variation around 0 time lag before and after learning. (Pos. mod. vs neg mod.: P < 0.0001; pos. mod. vs no mod.: P < 0.0001; KW test; n = 121; 29; 249 from 5 mice). **(C)** Time lag from 0 of the peaks or troughs of the CCG before and after learning for vLGN neurons positively modulated by plHVA activation. ($P < 1*10^{-31}$, Wilcoxon Signed-Rank Test; n = 121 cells). Statistical significance is indicated as *** P < 0.001.

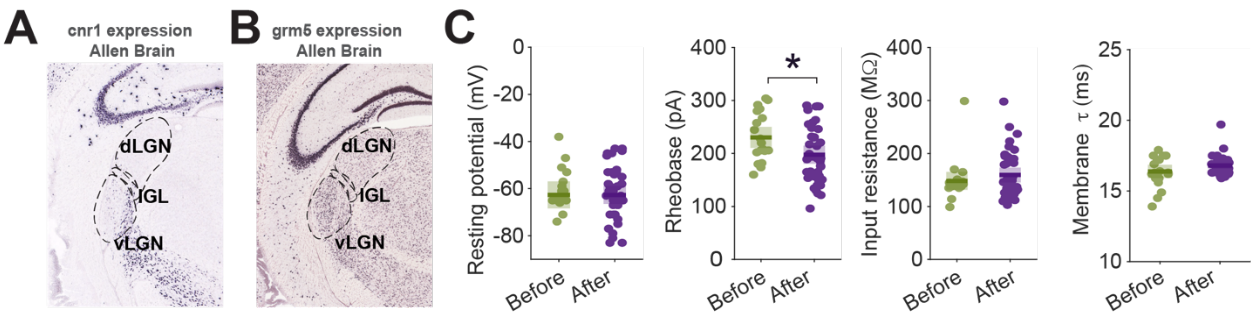


**Figure S7**. **Gene expression in vLGN and electrophysiological properties of vLGN neurons before and after learning.** (A) Gene expression of cnr1 (coding for the CB1 receptor) in a coronal slice of the vLGN, as assessed by in situ hybridization (ISH) in the Allen Brain Atlas (Seattle, WA, USA) (44). Note that CB1 receptors appear medial in vLGN, potentially presynaptic to the $vLGN^{plHVA}$ cells (B) Same as A, but for the grm5 gene (coding for mGluR5). (C) Boxplots for resting membrane potential, rheobase, input resistance, and membrane time constant recorded from $vLGN^{plHVA}$ neurons in brain slices prepared from mice before (green) and after learning (purple). ($P = 0.998$; $P = 0.024$; $P = 0.736$; $P=0.265$,KW test; $n = 20$ cells from 4 mice before learning, 39 cells from 5 mice after learning). Statistical significance is indicated as * $P < 0.05$. Dots indicate individual $vLGN^{plHVA}$ neurons.
